# Chromatin State Dynamics Confers Specific Therapeutic Strategies in Enhancer Subtypes of Colorectal Cancer

**DOI:** 10.1101/2020.09.04.283838

**Authors:** Elias Orouji, Ayush T. Raman, Anand K. Singh, Alexey Sorokin, Emre Arslan, Archit K. Ghosh, Jonathan Schulz, Christopher J. Terranova, Ming Tang, Mayinuer Maitituoheti, S. Carson Callahan, Katarzyna J. Tomczak, Zhiqin Jiang, Jennifer S. Davis, Sukhen Ghosh, Hey Min Lee, Laura Reyes-Uribe, Kyle Chang, Yushua Liu, Huiqin Chen, Ali Azhdarnia, Jeffrey S. Morris, Eduardo Vilar, Kendra S. Carmon, Scott Kopetz, Kunal Rai

## Abstract

The extent and function of chromatin state aberrations during colorectal cancer (CRC) progression is not completely understood. Here, by comprehensive epigenomic characterization of 56 tumors, adenomas, and their matched normal tissues, we define the dynamics of chromatin states during the progression of colorectal cancer. H3K27ac-marked active enhancer state could distinguish between different stages of CRC progression. By epigenomic editing, we present evidence that gains of tumor-specific enhancers for crucial oncogenes, such as *ASCL2* and *FZD10*, was crucial for excessive proliferation. Consistently, combination of MEK plus bromodomain (BET) inhibition was found to have synergistic effects in CRC patient-derived xenograft (PDX) models. Probing inter-tumor heterogeneity, we identified four distinct enhancer subtypes (EpiC), three of which correlate well with previously defined transcriptomic subtypes (CMSs). Importantly, CMS2 can be divided into two EpiC subgroups with significant survival differences. Leveraging such correlation, we devised a combinatorial therapeutic strategy of enhancer-blocking bromodomain inhibitors with pathway-specific inhibitors (PARPi, EGFRi, and TGFβi) for three EPIC groups. Our data suggest that the dynamics of active enhancer underlies colorectal cancer progression and the patient-specific active enhancer patterns govern their unique gene expression patterns which can be leveraged for precision combination therapy.

## INTRODUCTION

Colorectal cancer (CRC) is the third most common cancer in the world. Most cases of CRC originate from premalignant lesions, mainly adenomas, which can progress into malignant adenocarcinomas(1). Genomic alterations in colorectal adenocarcinomas frequently occur in *APC* (83%), *TP53* (60%), *KRAS* (43%), and *BRAF* (10%) genes(2-6). However, targeted therapy for these mutational subgroups is not well established. This calls for a comprehensive approach to identify unique signatures derived from various molecular characteristics of tumors including genomic and epigenomic components which, along with clinical features, can be used to introduce clinically feasible targeted therapeutic strategies.

One such effort toward precision therapy was led by the CRC subtyping consortium, which identified four consensus molecular subtypes (CMSs) with distinct activation patterns of specific oncogenic signaling pathways using gene expression datasets(3). Briefly, these transcriptomic subgroups include the immune subtype (CMS1), canonical subtype (CMS2), metabolic subtype (CMS3), and mesenchymal subtype (CMS4)(7). Although distal *cis*-regulatory elements are known to play a central role in controlling transcription patterns, the bona fide contributions of these elements in defining CMSs are not understood. Importantly, epigenetic processes are reversible and can provide actionable targets for therapeutic development, such as bromodomain inhibitors (BRDi) that target super-enhancer elements(8). The benefits of targeting the CRC epigenome have already been demonstrated by associations of the CpG island methylator phenotype (CIMP) with BRAF mutations, for which efforts are underway to devise combinatorial strategies targeting methylation and the BRAF oncogene(3). Consistent with this finding, we and others have provided evidence that epigenomic alterations at the level of DNA methylation play crucial roles in CRC initiation(9) and oncogenic function(10).

The current literature on alterations of enhancer elements in CRC is scarce, with only a handful of studies(11-13) with small sample sizes that profiled primary tumors. Furthermore, a comprehensive understanding of other chromatin state alterations and their functional contribution to tumor progression is mostly lacking. Therefore, it is of immense importance to define the relationship between chromatin aberrations and genomic and transcriptomic features in tumors.

## RESULTS

To characterize the epigenomic makeup of CRC, we first generated ChIP-seq (chromatin immunoprecipitation followed by massive parallel sequencing)-based profiles for six histone marks including H3K4me1 and H3K27ac, associated with enhancer regions; H3K4me3, associated with promoter regions; H3K79me2, associated with transcribed regions; H3K9me3, associated with heterochromatin regions; and H3K27me3, associated with Polycomb repression in a total of 16 CRCs and 17 matching adjacent normal colonic mucosa samples (**Table S1**). With the use of the ChromHMM algorithm(14), we trained a 10-state, 12-state, and 15-state model (for samples having all the six marks; Table S1) consisting of combinatorial patterns of specific histone modifications known as *chromatin states*, which are linked with the transcriptional output of the associated genomic loci and regulate the recruitment of various components of the transcriptional machinery. We chose a 10-state model for in-depth investigation because it represented biologically interpretable combinatorial patterns that were not repetitive or ambiguous (**Fig. 1A and S1A**). We annotated these chromatin states based on the content of each state and its average relative distribution around known genomic features (**Fig. S1B**). To define the chromatin states that best differentiate between normal and tumors, we developed a statistical approach, termed ChromXplorer (see Methods; **Fig. S1C**), that utilizes the ChromDiff(15) framework for all genomic loci instead of a gene-centric approach. The ChromXplorer-based dimensionality reduction approach showed the “active enhancer” state, but not others, to be the most discriminatory state in all three investigated models (**Fig. 1B and Fig. S1D-S1F**). Analysis of singular histone marks indicated that the active enhancer mark, defined by H3K27ac, also led to clustering of the samples according to their phenotype (**Fig. S2A**), whereas some of the other histone marks such as H3K79me2, H3K9me3, and H3K27me3 showed weaker clustering patterns due to variations among tumor samples (**Fig. S2A**), suggesting *de novo* gain or loss of H3K27ac patterns as an important distinguishing feature of CRC.

**Fig. 1.**
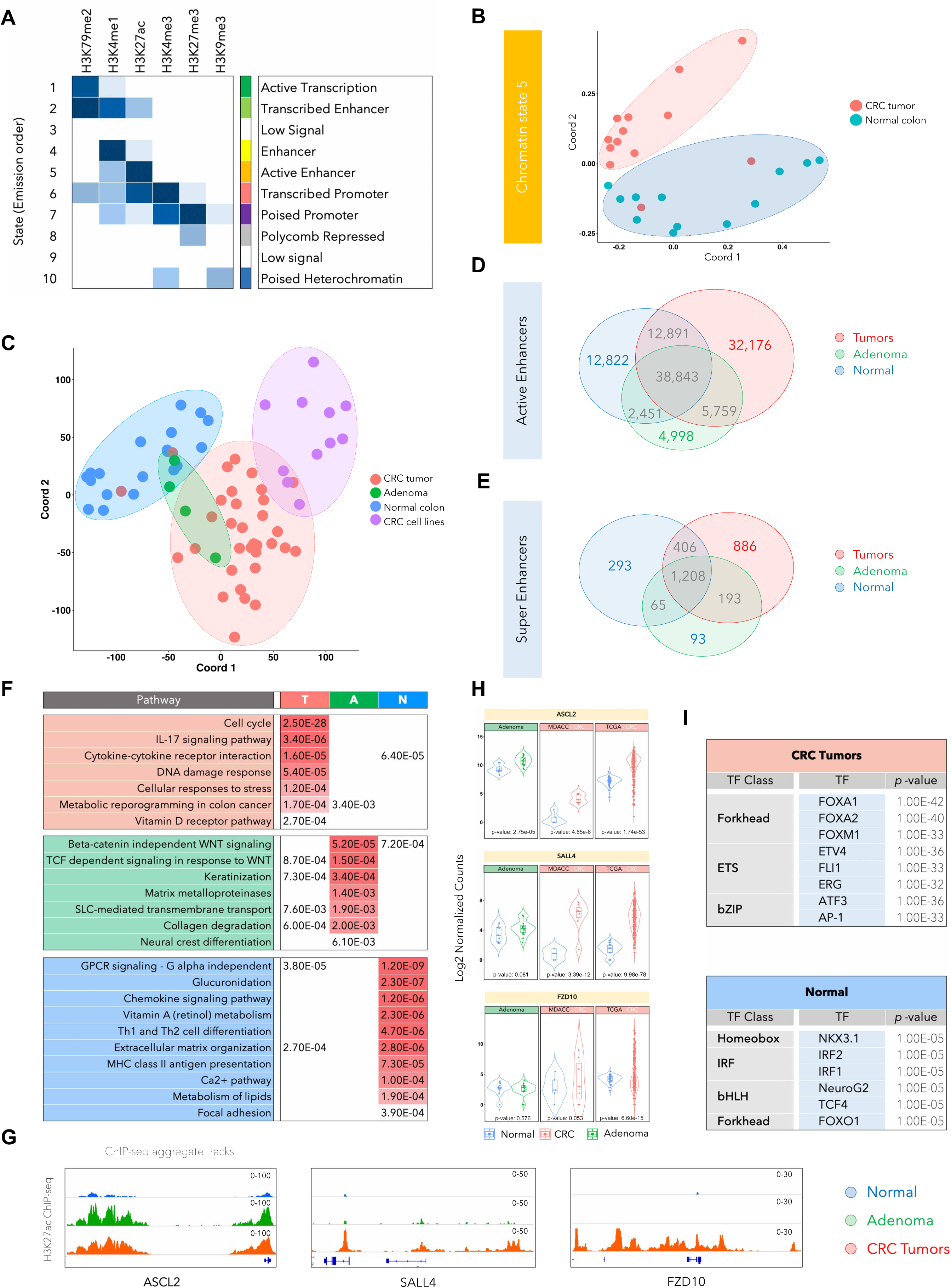
H3K27ac-identified global enhancer pattern could differentiate colorectal cancer from normal colon. A. Emission probabilities of ChromHMM-based 10-chromatin state calls, which is based on the global distribution of combinatorial patterns of 6 histone marks (H3K4me1, H3K27ac, H3K4me3, H3K27me3, H3K79me2, and H3K9me3) from 13 adenocarcinomas and 12 adjacent normal tissues. Each row is representative of a combinatorial chromatin state pattern, which is annotated on the right based on the constituent histone mark and neighborhood plot (Fig. S1B). Columns represent each histone mark. Colors represent the frequency of occurrence of that mark in the corresponding chromatin state on a scale of 0 (white) to 1 (blue). B. Multidimensional scaling (MDS) plot for chromatin state 5 between normal tissues (blue) and adenocarcinoma samples (red). C. MDS plot for global H3K27ac patterns in the adenomas (A), adenocarcinomas (T), adjacent normal colon (N) and cancer cell lines. Adenoma samples are spread over normal and tumor samples and cannot be distinguished with use of H3K27ac. Red, Green, Blue and Purple colors were used for Tumor, Adenoma, Normal Colon and CRC cell lines respectively. D. Venn diagram showing overlaps between total number of H3K27ac-defined active enhancer peaks in normal samples (blue), adenomas (green) and CRC tumors (red). E. Venn diagram showing overlaps between total number of super-enhancer peaks (called using ROSE) in normal samples (blue), adenomas (green) and CRC tumors (red). F. List of significantly enriched pathways in genes targeted by enhancers specific to CRC tumors (T), adenomas (A), and normal colon tissue (N). *q*-values are shown and are color-coded based on the level of significance. G. Integrative Genomics Viewer (IGV) images showing enrichment of H3K27ac peaks around ASCL2, SALL4 and FZD10 genes using aggregate ChIP-seq profiles of normal colon, adenoma and CRC tumors. H. Violin plots showing mRNA expression levels for ASCL2, SALL4 and FZD10 in adenomas, the adenocarcinoma cohort used in this study (labeled as MDACC) and TCGA adenocarcinoma cohort. The reported p-values were calculated using the *DESeq2* (60). I. List of enriched transcription factor (TF) motifs in CRC- or normal tissue-specific active enhancers. Motifs are identified using HOMER(25) on the basis of the shared H3K27ac peak in each group.

In light of this, we expanded the cohort by profiling the H3K27ac mark in additional colorectal tumors (n = 17), adenomas (n = 4), normal crypts (n = 4), and cancer cell lines (n = 11) to characterize the enhancer patterns through the various stages of progression. Interestingly, distinct enhancer patterns appeared to define each progression stage, suggesting that these enhancer patterns may evolve as the disease progresses (**Fig. 1C**). Furthermore, the established CRC cell lines formed a separate cluster with three cell lines that partially overlapped with colorectal adenocarcinoma tumors (**Fig. 1C**). Similarly, computation of correlation matrix identified a higher similarity within normal samples as opposed to tumor samples which segregated into two subgroups, indicative of greater heterogeneity between adenocarcinoma specimens (**Fig. S2B**).

To identify recurrent cancer-specific peaks, shared H3K27ac peaks in all tumor samples were summed up and overlapped with shared normal or shared adenoma enhancers. Our analysis identified 32,176 tumor-specific (T) enhancers, 12,822 normal-specific (N) enhancers, and 4,998 adenoma-specific (A) enhancers (**Fig. 1D**). Notably, 5,759 enhancers were present in both adenomas and adenocarcinomas, but not in normal tissue, whereas a much higher number (32,176) were specific to adenocarcinomas only, thereby, suggesting that several enhancers are gained in *de novo* manner in the adenocarcinoma stage. Furthermore, the average intensity of enhancer peaks was found to progressively increase in adenomas and colorectal tumors (**Fig. S2C**). Similar to typical enhancers, we found super-enhancer elements (SE), that are thought to promote oncogene expression, to be upregulated in adenocarcinomas (**Fig. 1E**). We identified 886 adenocarcinoma-specific, 93 adenoma-specific, and 193 shared super-enhancers between adenomas and adenocarcinomas using ROSE algorithm(16) (**Fig. 1E**).

Active enhancers specific to each disease progression stage marked important driver genes that were identified based on the presence of enhancers in their proximal regulatory regions as previously defined(17,18) (**Fig. 1F and Table S2**). For example, regulators and mediators of the cell cycle, DNA damage, stress and cytokine and metabolism harbored gains in a tumor-specific manner. Conversely, enhancers for regulators of T-cell differentiation, MHC class II antigen presentation, vitamin A metabolism, chemokine signaling, and focal adhesion, among others, were specifically present only in normal tissues and were lost during progression. Enhancers common to premalignant adenomas and tumors were enriched around regulators of cell fate, collagen degradation, metabolic transport, and WNT signaling (**Fig. 1F**). We noted that the enrichment of enhancers/super-enhancers in the proximity of key WNT signaling regulators such as ASCL2, SLL4, FZD10, CTNNB1 (β-catenin), AXIN2, TCF7L1 (TCF3), TCF7L2 (TCF4), and a subset of these correlated with their progressive increase in gene expression (**Fig. 1G-H, Fig. S3A-C, and Table S2**). This suggests that the genes identified to harbor active enhancers or super-enhancers near their vicinity or in regions interacting with the gene body/promoters are likely to contribute to the carcinogenic properties. Enhancers are epigenetic platforms for binding of various transcription factors (TFs), and motif enrichment analysis has suggested that specific classes of TFs such as forkhead (FOXA1, FOXA2, and FOXM1), ETS (ETV4, ERG, and FLI1), and bZIP (ATF3 and AP-1) are overrepresented in the tumor-specific active enhancers (**Fig. 1I and Fig. S3D**). Importantly, homeobox (NKX3.1), IRF (IRF1, IRF2), bHLH (NeuroG2, TCF4), and FOXO1 motifs were enriched in active enhancers that were lost during tumor progression (**Fig. 1I and Fig. S3D**).

### Inactivation of tumor-specific enhancers leads to the transcriptional and phenotypical switch

To test the functional role of some of the specifically activated tumor-specific enhancers, we first ranked the top differentially activated enhancers (**Fig. 2A and Table S3;** see Methods). We chose to functionally test some of the top enhancers from this list that were located in the vicinity of two WNT signaling genes: *ASCL2* and *FZD10. ASCL2* is a known driver of intestinal stem cell phenotype(19) and a predicted oncogene in CRCs(20) whereas, *FZD10* is a transmembrane protein that acts as a WNT receptor driving downstream signaling, which is a key signaling event in CRCs(7). Moreover, established CRC cell lines harbored a significant percent of tumor-specific H3K27ac peaks (**Fig. S4A**). We used dCas9-KRAB and specific gRNAs to suppress the activity of target enhancer loci in SW480 cells, a well-characterized model with mutations in *APC, KRAS*, and *TP53*^20^. For ASCL2, we chose to focus on a distal super-enhancer that was present in the adenomas and adenocarcinomas, but not normal tissues (**Fig. 2B, 1H, and S3A**). HiChIP and 3C experiments confirmed the direct interaction of this super-enhancer with the ASCL2 gene promoter (**Fig. 2B and S4B**). We noted that all three gRNAs targeting this distal super-enhancer region were able to suppress the H3K27ac signal in their respective target regions (**Fig. 2C**). These gRNAs also downregulated gene expression (**Fig. 2D**) and downstream proliferation of SW480 cells, suggesting that the distal super-enhancer acts as a functional driver event in this cell line (**Fig. 2E**). For the FZD10 locus, we noted that gRNAs targeting E1, E5, and E6 regions reduced the H3K27ac signal which was consistent with the observations from HiChIP experiments showing the interaction of these enhancers with the promoter of this gene (**Fig. S4C-4D**). Epigenetic suppression of these enhancers also downregulated gene expression and decreased cell proliferation, suggesting multi-enhancer regulation of FZD10 transcription (**Fig. S4E-4F**) and the contribution of this locus to the proliferative properties of CRC cells (**Fig. S4F**).

**Fig. 2.**
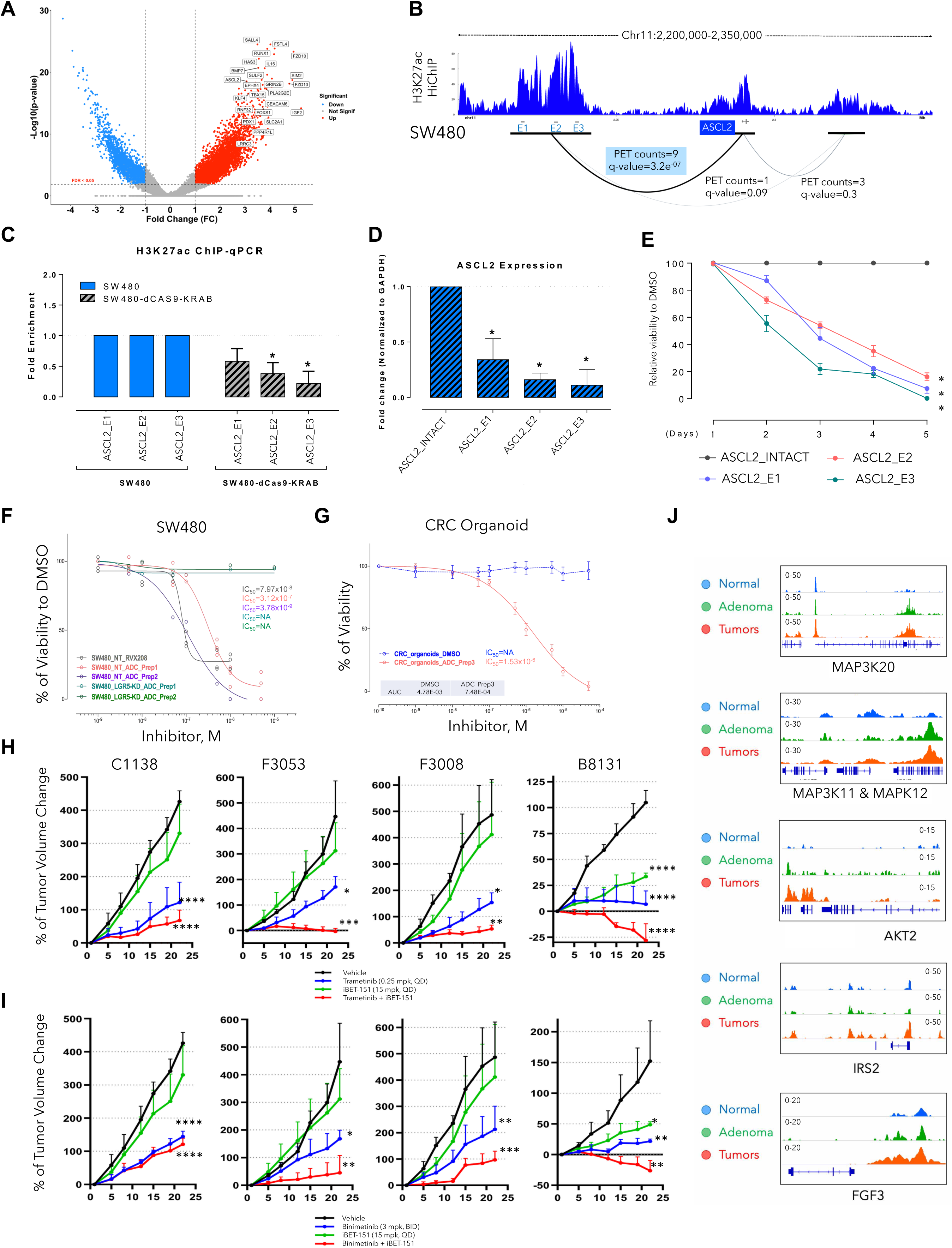
Inactivation of top tumor-specific enhancers leads to transcriptional downregulation and loss of cell proliferation. A. Volcano plot showing top CRC-enriched typical enhancers ranked based on fold-change and p-value. Top *de no*vo gained enhancers among tumor (n = 33) as compared to adjacent normal (n = 15) samples are annotated according to their associated genes. Thresholds of log2FC > 1 and FDR < 0.05 are used to annotate *de novo* enhancer peaks. ASCL2 and FZD10 enhancers are among this subset of highly enriched regions. There are two top ranked enhancer peaks around FZD10 gene. B. Interaction map of active enhancers (E) with the gene promoter (P) around the ASCL2 locus as determined in SW480 cell line by HiChIP. Interactions were observed in a super-enhancer (E1-E3) region located ∼70kb upstream of ASCL2 promoter (PET counts=9, *q*-value=3.2e^-07^). These regions are among the highly enriched tumor-specific enhancers and were present in multiple CRC cell lines (Figure S4A). C. Bar graph showing relative enrichment of H3K27ac as derived from a ChIP-qPCR experiment on E1-E3 for ASCL2 super-enhancer (from panel B) in SW480 parental or modified cells harboring gRNAs to these enhancers. Y-axis represents fold change in intensity of H3K27ac signal for each enhancer in gRNA harboring cell in comparison to the parental cell. Asterisk * denotes p-value < 0.05. D. Bar graph showing relative ASCL2 mRNA expression in E1-E3 gRNAs for ASCL2 super-enhancer harboring SW480 cells in comparison to parental cells (ASCL2_INTACT). E. Growth curve showing relative viability of SW480 cells harboring E1-E3 for ASCL2 super-enhancer in comparison to parental cell (ASCL2_INTACT) showing lower rates of proliferation in cells with ASCL2-inactivated enhancers. F. Growth curves for LGR5-silenced SW480 cells (SW480^LGR5-KD^) in comparison to the parental cells upon treatment with increasing concentrations of two preparations of antibody-drug conjugate (ADC) with bromodomain inhibitor (RVX208) conjugated to LGR5 antibody (ADC-Prep1 and ADC-Prep2) and RVX-208 alone. Y-axis shows percent viability of treated cells in comparison to control (DMSO) treated cells. G. Growth curves for a patient-derived CRC organoid upon treatment with LGR5-RVX208 conjugate (ADC). Y-axis shows percent viability of treated cells in comparison to control (DMSO) treated cells. H. Tumor volume curves for C1138, F3053, F3008, and B8131 on treatment with MEK inhibitors (trametinib), BRD inhibitor (i-BET151), and a combination of MEK and BRD inhibitors (trametinib + i-BET151) along with the control vehicle group (n = 3-5 for each arm). P-values represent pairwise *t* test comparison between the experimental arm to vehicle treatment. * = p < 0.05; ** = p < 0.01; *** = p < 0.001; **** = p < 0.0001. Please refer to **Figure S6B** for all pair-wise comparisons of the p-values. I. Tumor volume curves for C1138, F3053, F3008, and B8131 on treatment with MEK inhibitor (binimetinib), BRD inhibitor (i-BET151), and a combination of MEK and BRD inhibitors (binimetinib + iBET151) along with the control vehicle group (n = 3-5 for each arm). P-values represent pairwise *t* test comparison between the experimental arm to vehicle treatment. * = p < 0.05; ** = p < 0.01; *** = p < 0.001; **** = p < 0.0001. Please refer to **Figure S6B** for all pair-wise comparisons of the p-values. J. IGV images showing enrichment of H3K27ac peaks around MAP3K20, MAP3K11, MAPK12, AKT2, IRS2, and FGF3 genes using aggregate ChIP-seq profiles of normal colon, adenoma and CRC tumors

Although these studies provide evidence for the functionality of individual enhancers, a holistic approach would be to block many such enhancers using BRDi, which are being actively tested in clinical trials in solid tumors. Since ASCL2 and WNT pathways have established roles in regulating stem cell function in both normal and cancer tissues(7,19,20), we asked whether targeting enhancer-blocking agents to stem cells, using the LGR5 antibody, would result in suppressing the growth of the cells. Hence, we synthesized an antibody-drug conjugate (ADC) by using LGR5 antibody and the RVX208 which is a BRDi in phase 3 trials for cardiovascular indications(21). Indeed, LGR5-RVX208 (**Fig. S5A**) ADCs specifically blocked the growth of SW480 cells or patient-derived human colon organoids in an LGR5 expression–dependent fashion (**Fig. 2F-2G and Fig. S5B-E**). These studies established that some of the activated enhancers in tumors may play functional roles in imparting pro-tumorigenic properties to CRC.

Treatment of four different patient-derived xenograft (PDX) models (C1138, F3053, F3008, and B8131) with BRDi (iBET-151) caused strong tumor reduction in one model but showed no significant difference in 3 of 4 models, suggesting a context-dependent impact of enhancer blockade (**Fig. 2H-I)**. Since chromatin aberrations are believed to be a permissive event for activation of pro-tumorigenic transcriptional signature upon oncogenic signaling, we reasoned that blockade of enhancers may be more effective in combination with inhibition of key oncogenic driver signaling pathways. We noted that a number of upstream regulators of the MEK pathway, including *EGFR, NRAS, KRAS, FGFR3, MAPK3K1, MAPK11, MAPK12, MAP3K20, MAP3K10, AKT2, IRS2, FGF1, FGF3, FGF19*, and *FGF21*, gained enhancers in colorectal tumors in comparison to adjacent normal tissues or adenomas (**Fig. 2J and S6A**). Therefore, we analyzed whether enhancer-blocking BRDi may synergize with MEKi, which are actively being used in CRC clinical trials, to enhance therapeutic efficacy. Indeed, the treatment of four different PDX models (C1138, F3053, F3008, and B8131) showed significant synergy between MEKi (trametinib or binimetinib) and BRDi in 3 of these 4 models (**Fig. 2H-I and S6B)**. Although monotherapy with MEKi showed moderate activity on tumor growth in all of the models, significant tumor regression was noted in B8131, and two models (F3052 and 3008) showed slower growth in combinatorial treatment with BRDi. These data highlighted the differential responses of various PDX models with BRDi and suggested that BRDi plus MEKi could be a successful therapeutic strategy in a subset of CRC patients.

### Distinct enhancer-driven EPIgenome-based Classification (EpiC) clusters of CRC

As exemplified by the MEKi + BRDi combination treatment results, all CRC tumors do not respond to the blanket therapies equally. This motivated us to perform molecular characterization of CRC tumors based on their epigenomic content, which can be potentially useful for patient stratification and subsequent assignment of each subtype to particular combination treatment with epigenetic compounds.

We classified adenomas and adenocarcinomas by using a consensus non-negative matrix factorization (NMF)-based clustering method that uses the enriched H3K27ac peaks. Based on the cophenetic and dispersion scores, we identified four distinct clusters of tumors (**Fig. 3A, S7A S7B, and Table S4**), hereafter referred to as <u>EPI</u>genome-based <u>C</u>lassification (EpiC) clusters: EpiC1, EpiC2, EpiC3, and EpiC4. The average intensity of enhancer peaks within these clusters varied such that EpiC2 and EpiC4 enhancers had significantly higher peaks than the other two clusters had (**Fig. S7C-S7D**). EpiC3 clusters had the least overall peak intensity (**Fig. S7C-S7D**). However, differential regions across these subtypes were found to be specific to particular genes (**Fig. S8A**).

**Fig. 3.**
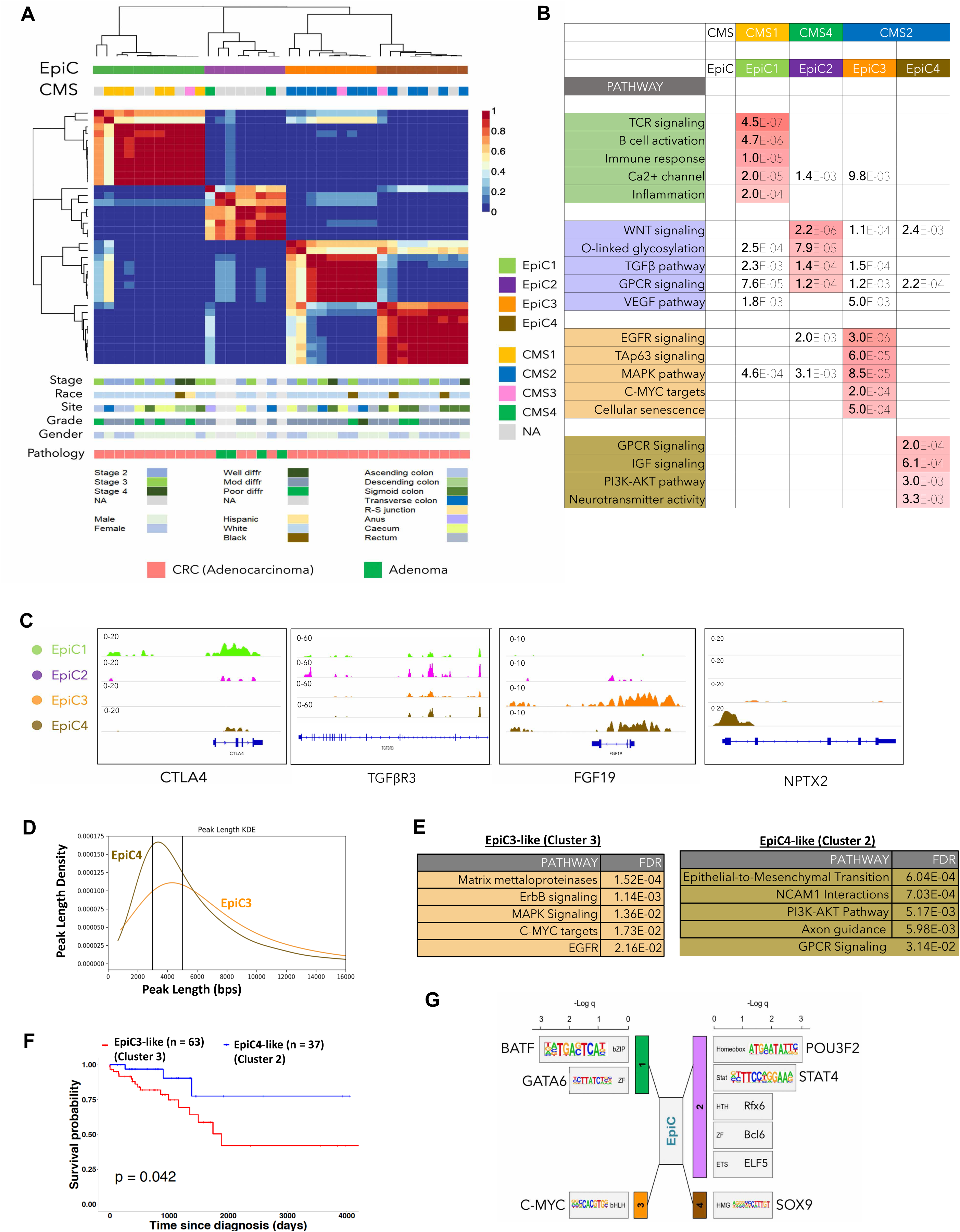
Reclassification of colorectal cancer tumors based on their global enhancer distribution leads to identification of enhancer-based subtypes. A. Non-negative matrix factorization (NMF) clustering of CRC adenocarcinoma (n = 33) and adenoma (n = 4) samples identified four EPIgenome-based <u>C</u>lassification (EpiC) clusters of CRC, shown in the color-coded matrix on the top panel. Consensus molecular subtype (CMS) classification of each tumor sample is identified by using a 498 gene–based random forest approach and is overlaid with the EpiC clusters. Clinical data corresponding to the tumor samples in each cluster, including tumor stage, site, pathological grade, race, and gender of the patients, are provided in the lower panels. B. Table showing enriched major signaling pathways for genes targeted by EpiC-specific enhancers using GREAT(17). False Discovery Rate (FDR) values are color-coded. C. IGV images showing enrichment of H3K27ac peaks around CTLA4 for EpiC1, TGFβR3 in EpiC2, FGF19 in EpiC3, and NPTX2 in EpiC4 using aggregate ChIP-seq profiles of EpiC1, EpiC2, EpiC3 and EpiC4 group tumors. D. KDE (Kernel Density Estimate) plot with peak lengths (x-axis) and densities (y-axis) of EpiC3 and EpiC3-specific enhancers. Peaks with length more than 5kbs were annotated as Broad whereas those with < 3kb in length were called as Sharp. E. Pathway analysis for genes in Clusters 2 and 3 from NMF clustering (see Figure S9) of 115 CMS2 TCGA CRC tumors that overlap with EpiC3 and EpiC4-unique genes (LogFC >0.5 and p-value < 0.05). Based on the pathway enrichment, Cluster 2 was annotated as “EpiC4-like” whereas Cluster 3 was annotated as “EpiC3-like”. F. Kaplan-Meier plot of survival of CMS2 samples in the TCGA, comparing EpiC4-like patients (n = 37) and EpiC3-like patients (n = 63). The log-rank (Mantel-Cox) p-value is shown for the difference in survival. G. Motif enrichment analysis for enhancers unique to different EpiC groups using HOMER (25).

Previously, four CMSs had been defined on the basis of transcriptomic signatures of more than four thousand CRC samples (3) as the CMS1 (MSI immune subtype), CMS2 (canonical subtype), CMS3 (metabolic subtype), and CMS4 (mesenchymal subtype). We classified the tumors used in our study into these CMSs, for which we had gene expression data, by using the CMS classifier(22), and based on the corresponding trained model(3), we computed their EpiC subtype. Interestingly, we determined that epigenomic clusters (EpiCs) significantly overlap their assigned CMSs (**Fig. 3A**). All of the samples classified as CMS1 (immune) subtype overlapped with the EpiC1 subtype. Although only two patients had been assigned to the CMS4 (mesenchymal) subtype, strikingly, all four adenoma samples along with these two patients clustered as the EpiC2 subtype. It has been shown that adenoma lesions have mesenchymal properties and may progress to adenocarcinoma due to aberrations in the Ras signaling pathway (23). Moreover, consensus NMF clustering divided the canonical subtype or CMS2 into two novel subsets of epigenomic CRC clusters, EpiC3 and EpiC4 (**Fig. 3A**). Of note, the metabolic subtype or CMS3 is not detectable with the use of enhancer-based epigenomic clustering and is scattered in multiple subtypes.

Interestingly, pathway analysis based on the gene targets of enhancers (see Methods) that are specifically enriched in each EpiC subtype showed similarity to those known to be CMS-specific (3) (**Fig. 3B**). Enhancers specific to EpiC1, which overlaps significantly with CMS1, were in the proximity of genes regulating immune response, B- and T-cell activation, and inflammation. EpiC2, which overlaps with CMS4 and interestingly contains all adenoma samples, regulates the TGFβ, WNT, and VEGF pathways (**Fig. 3B**). CMS2 CRC tumors were distributed into two epigenetic subtypes EpiC3 and EpiC4. EpiC3 enhancers enriched on targets of c-MYC, EGFR, MAPK, and TAp63 signaling, whereas EpiC4 enhancers activated genes involved in GPCR, IGF, PI3K-AKT and neurotransmitter signaling (**Fig. 3B and Table S5**). Examples of genes near the enhancers in each EpiC category are shown in **Fig. 3C** and **Fig. S8**.

We also noted that EpiC3 enhancers were more spread (or “broad”) in nature whereas EpiC4 enhancers were narrower in their width (or “Sharp”) (**Fig. 3D**). Systematic analysis of broad (>5kb) and sharp (<3kb) peaks (see Methods) showed enrichment of broad peaks in EpiC3 versus sharp peaks in EpiC4 enhancers (**Fig. 3D and S7E**).

To determine if the observed dichotomy of CMS2 tumors into EpiC3 and EpiC4 in our cohort expanded to a larger set of CRC samples, we examined the clustering of CMS2-designated TCGA tumors (24) based on the gene targets of EpiC3 and EpiC4 enhancers. NMF clustering of 115 CMS2 tumors (see Methods) identified 3 clusters (**Fig. S9A and Table S6**). Overlap of EpiC4 enhancer target genes with Cluster2-specific genes showed enrichment of EpiC4-specific pathways such as GPCR, PI3K pathway and NCAM mediated neurotransmitter signaling (including NPTX2 – a differentially expressed gene between EpiC3 and EpiC4) (**Fig. S9B-D**). Overlap of EpiC3 enhancer target genes with Cluster3-specific genes showed enrichment of EpiC3-specific pathways such as EGFR signaling, MAPK, and c-MYC targets (**Fig. 3E and S9B-C**). On the contrary, EpiC3-target genes overlap with Cluster3-specific genes or EpiC4-target genes overlap with Cluster2-specific genes did not enrich in any pathways specific for EpiC3 or EpiC4 (**Fig. S9C)**. Cluster 1 specific genes were mostly enriched in immune pathways suggesting likely separation due to low tumor content (**Fig. S9C)**. Therefore, we designated Cluster2 (n = 37) as ‘EpiC4-like’ and Cluster3 (n = 63) as ‘EpiC3-like’. Importantly, EpiC3-like tumors showed poorer survival in comparison to EpiC4-like tumors (**Fig. 3F)**, thus providing clinical significance for enhancer-based classification of CRC tumors.

To identify potential regulators of the EpiC-specific features, we performed motif analyses of enhancers specific to each EpiC subtype using HOMER (25). Interestingly, distinct TF classes were enriched in each of the EpiC subtypes. bZIP (BATF) and ZF (GATA6) type TFs were enriched in EpiC1, and homeobox (BRN2 or POU3F2) and Stats (STAT4) were enriched in EpiC2; c-MYC (bHLH) was enriched in EpiC3, and HMGs (SOX9) were enriched in EpiC4 (**Fig. 3G**). Consistently, we noted that expression of TFs (except for POU3F2) was higher, specifically in the respective CMSs in TCGA samples that corresponded with EpiC subtypes (**Fig. S9E**). Importantly, previous studies have implicated c-MYC(26-29), GATA6(30-32), SOX9(33-35), and STAT4(36,37) in CRC progression. BRN2/POU3F2 has oncogenic roles in melanoma(38,39), and BATF2, a BATF family member, has been implicated in CRC(40). These data suggest that the potential origin of different EpiC subtypes may be due to the pioneering activity of TFs belonging to different classes of TF families.

### Combinatorial treatments for newly identified EpiCs

The CMS classification is being tested as a prognostic indicator for use in targeted therapy in CRC. For example, the CMS1 and CMS4 subtype cell lines are more responsive to HSP90 inhibitor, whereas the CMS2 subtype shows more sensitivity to WNTi and EGFRi(41,42). The CMS3 subtype is enriched in metabolic pathways and RAS mutations, but therapeutic vulnerabilities are unknown(3).

Exploring the clinical implications of our findings, we used a panel of 11 signaling pathway inhibitors that are known to be involved in the CRC tumorigenesis and whose key regulators showed differential enhancer enrichments in their vicinity in different epigenomic subtypes of CRC. Therefore, we treated six CRC cell lines with the signaling pathway compounds along with a panel of 8 known epigenetic inhibitors to treat cell lines from different CMSs, previously annotated with use of a CMS classifier **(Fig. S10A, S10B, S11, S12, S13A, and S13B**)(41,42). We further analyzed both IC_50_ and AUC indices to choose compounds with the best potency as well as the highest efficiency (**Fig. 4A-4C, S10B, S13A**). We used a combination of one signaling pathway inhibitor along with the most efficient epigenetic inhibitor according to the generated drug matrix in distinct CMSs. Based on the analysis of monotherapies, we tested combinations of BRDi + PARPi for CMS1/EpiC1, BRDi + EGFRi for CMS2/EpiC3-4, and BRDi + TGFβi for CMS4/EpiC2 and found the combinations to perform better in reducing cell growth *in vitro* (**Fig. 4D-4F and Figs. S13B and S14A**). Importantly, we observed drastically enhanced activities of these combinations in the *in vivo* experiments carried out in nude mice, compared with the results of experiments using monotherapies (**Fig. 4G-4I**).

**Fig. 4.**
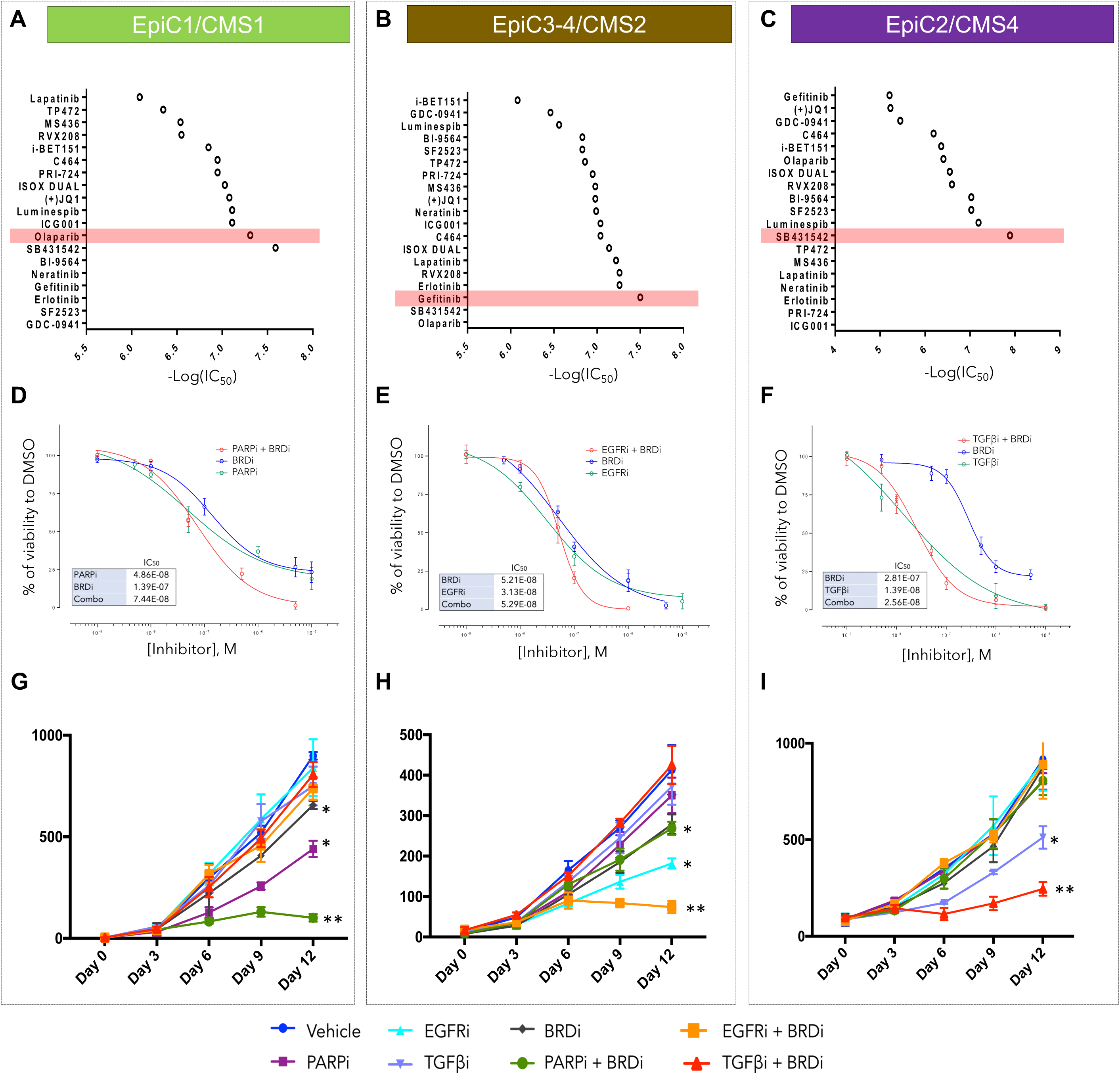
Correlation between CRC molecular subtypes and EpiCs can predict combinatorial treatments with high clinical significance. **A-C**. Summary of IC_50_ values for 19 small molecule inhibitors (Y-axis) in cell lines with similar expression features to CMS1/EPIC1 (HCT116 or SW480) (A), CMS2/EPIC3 or EPIC4 (T84) (B), and CMS4/EPIC2 (SW620) (C). -Log(IC_50_) values are demonstrated on X-axis for each of the compounds. Drugs with higher -Log(IC_50_) are highlighted and were further used in combinatorial experiments. **D-F**. Growth curves showing responses of CMS-specific CRC cell lines for the drug combinations identified based on the IC_50_ and AUC indices (panels A-C and Figures S8-S11): CMS1 lines with PARPi (olaparib) + BRDi (iBET-151) (D), CMS2 with EGFRi (gefitinib) + BRDi (RVX-208) (E), and CMS4 with TGFβi (SB431542) + BRDi (ISOX-DUAL) (F). **G-I**. Tumor volume curves for xenografts in NUDE mice generated from transplantation of EpiC1/CMS1 (G), EpiC3-4/CMS2 (H), and EpiC2/CMS4 (I) cell lines upon treatment with inhibitors of EGFR (gefitinib, 100mg/kg), PARP (olaparib, 50mg/kg), TGF-β (SB431542, 10mg/kg), BRDi (i-BET151, 15mg/kg), or the combination of EGFRi + BRDi, the combination of PARPi + BRDi, and the combination of TGF-βi + BRDi along with the control vehicle group. Mice were treated every other day. Best treatment response was observed in the PARPi + BRDi combination in EpiC1/CMS1, EGFRi + BRDi combination in EpiC3-4/CMS2, and TGF-βi + BRDi combination in EpiC2/CMS4 lines. * shows p-values <0.05, and ** shows p-values <0.01.

Importantly, these combinatorial therapies demonstrated specific activity to the corresponding CMS/EpiC and had a minor impact on other CMS/EpiC subtypes. Specifically, BRDi (iBET-151, 15mg/kg) + PARPi (Olaparib, 50mg/kg) showed more significant impact on CMS1/EpiC1 cells than CMS2/EpiC3-4 or CMS4/EpiC2 did; similarly, BRDi (iBET-151, 15mg/kg) + EGFRi (Gefitinib, 100mg/kg) impacted CMS2/EpiC3-4 cells specifically, and BRDi (iBET-151, 15mg/kg) + TGFβi (SB431542, 10mg/kg) reduced tumor growth of CMS4/EpiC2 cells specifically (**Fig. 4G-4I**). Notably, we did not observe much loss in tumor weight upon treatment with these drug combinations suggesting little or no side effects of these therapies (**Fig. S14B)**.

## DISCUSSION

Our study provides the most comprehensive data on basic epigenetic elements, in line with definitions from Roadmap/ENCODE, for human CRC, pre-malignant adenomas, widely used cell lines, and normal crypts/tissues. Importantly, these tissues/cell lines have associated genomic, transcriptomic, drug-response, and clinical datasets available. Thus, these collective epigenomic maps elucidate the epigenomic makeup of CRC and are an essential resource, along with the previous datasets(11-13), for the community to generate hypotheses regarding the roles of the epigenome in CRC progression. Based on this comprehensive resource, our unbiased chromatin state analyses identified the H3K27ac marked active enhancers are the most discriminatory epigenetic elements between the various histopathological stages of CRCs. Consistent with this, the number of enhancer peaks that differentiate adenomas, normal tissues, and tumors is very high, suggesting that enhancer activation is likely a major epigenetic aberration in CRC. We suggest that enhancer aberrations are a critical component of a permissive chromatin environment in cancer cells that allow oncogenic events/pathways to drive excessive tumor growth. In addition to the active enhancer state, we also noted the heterochromatin state (H3K9me3) to differentiate between tumors and normal tissues. This will need to be further examined for causative effects.

Most importantly, our work identifies heterogeneity in enhancer patterns between CRC patients and for the first time identifies epigenomic subtypes of CRC. We demonstrate that the EpiC subtypes correlate highly with the expression phenotype (CMS), in line with previous observations made in the pan-cancer TCGA ATAC-seq study(43), except that the CMS2 subtype was bifurcated into two distinct EpiC subtypes, EpiC3 and EpiC4. This is an interesting observation since we hypothesize that EpiC3 is likely driven by a c-MYC–regulated transcriptional program, whereas EpiC4 is driven by SOX9 and had exclusive activation of IGF and PI3K-AKT signaling. In addition, only EpiC4 showed activation of enhancers around important neuronal genes (**Figure 3C and S8**), such as NPTX2(44), which could be either due to neural invasion in this category of tumors(45) or epigenetic derepression of neuronal features. We suggest that inhibitors that target the c-MYC program could be more useful in EpiC3 subtypes, whereas SOX9 may be a target in EpiC4 subtypes. Importantly, the two TCGA subsets with EpiC3 and EpiC4 like features showed significant difference in survival, hence providing a clinical significance of this bifurcation of the CMS2 subtype. How these EpiC-specific enhancer patterns are set-up and how they cooperate with genetic events for the transformation of normal cells into cancers with unique transcriptional signatures need to be studied in the future.

We identified superenhancers surrounding ASCL2 and FZD10 to be necessary for colorectal cancer cell growth in vitro (Figures 2 and S4). This is interesting in that it provides evidence for enhancer/superenhancer elements as being targets for therapeutics in the future, potentially with the evolution of CRISPR-based therapies. It is also imaginable that eRNAs made from such superenhancers may be targets for RNA-targeted therapeutics such as antisense oligonucleotides (ASOs)(46). The advances in 3D epigenome studies, such as our HiChIP data, also suggest that one enhancer may target many genes and hence could prove to be a better therapeutic target than the presumed gene target itself. We found over >30,000 such enhancers to be uniquely activated in colorectal adenocarcinomas. Future studies will be needed to identify functional enhancers such as those surrounding ASCL2 and FZD10. Finally, our data suggest that combination of bromodomain inhibitors as an umbrella approach for blockade permissive enhancer states along with specific driver signaling pathway inhibitors could be used in defined patient CRC populations, hence providing a paradigm for epigenome-focused personalized therapeutic strategy. This is an important observation as it suggests patient-stratification based on enhancer signatures as well as the use of epigenetic therapy in a patient-centric manner.

## METHODS

### Cell culture

Colorectal cancer cell lines were obtained from the MDACC cell bank. CRC cell lines including SW480, SW837, SW620, SW403, CW2, HCT15, HCT116, DLD1, SNUC5, CW2, and RKO were maintained in L-15 media with 10% FBS in tissue-culture conditions without CO_2_.

### CRC organoid culture

Tumor tissue dissociation was performed by using the MACS dissociator (Milteny Biotech). Sterile 24-well plates were pre-warmed in the 37°C incubator for ∼3 hours. Matrigel (Corning, Cat# 356231) was thawed on ice before use. Cells were resuspended in Matrigel at a concentration of 15-25k cells per 30 μl in each well. Matrigel-cell suspension was pipetted to the center of each well, and after 1-2 min at room temperature, the plate was transferred to a 37°C, 5% CO_2_ incubator for 5-10 min, followed by the addition of 500 μl of complete growth medium. The medium was replaced every 2-3 days and passaged every 7 days. The base medium was Advanced DMEM/F-12 (Life Technologies, Gibco) supplemented with B27, 2 mM (Life Technologies, Cat# 17504-044), GlutaMAX-1 (Life Technologies, Gibco), 10 mM HEPES (Life Technologies, Gibco), 1.25 mM N-acetyl cysteine (Sigma-Aldrich, Cat# A9165), 10 nM [Leu15]-Gastrin 1 (Sigma-Aldrich), and Pen/Strep (Life Technologies, Gibco). Complete growth medium consisted of base medium plus 50 ng/ml EGF (Life Technologies, Cat# PMG8043), 100 ng/ml Noggin, 0.5 μM A 83-01 (Tocris Bioscience), 10 μM Y-27632 (Sigma-Aldrich), 40% Wnt-3a, 10% R-spondin 1, 10 μM SB202190 (Sigma-Aldrich), and 5 μl of CHIR 99021 (Tocris Bioscience).

### Tissue samples and tissue preparation

All adenocarcinoma and normal tissue samples were obtained from the Division of Gastrointestinal Medical Oncology patients’ databank as frozen samples and were processed according to our modified protocol(47). Crypts were isolated from endoscopic biopsies of Familial Adenomatous Polyposis (FAP) patients that provided written informed consent to an IRB-approved protocol (MD Anderson Cancer Center #PA12-0327) following previously reported protocol (48). Detailed clinical and pathology characteristics of all samples are provided in Table S1. Crosslinked tissue samples were stored in a freezer at –80 °C before ChIP-seq experiments.

### Chromatin immunoprecipitation followed by sequencing (ChIP-seq)

Chromatin immunoprecipitation followed by deep sequencing was performed as described previously(49). Chromatin shearing conditions were optimized for colon cells. The following antibodies were used in this procedure; H3K4me1 (ab8895), H3K27ac (ab4729), H3K4me3 (ab8580), H3K79me2 (ab3594), and H3K27me3 (ab6002). Generated ChIP-seq data underwent quality-control measures and was further processed by pyflow-ChIPseq, a snakemake-based ChIPseq pipeline. Briefly, raw reads were mapped to human reference genome hg19 by using bowtie1, and duplicated reads were removed, retaining only uniquely mapped reads. RPKM-normalized bigwigs were generated by deep tools, and the tracks were visualized with the use of an Integrative Genomics Viewer (IGV).

### ChIP-seq analyses

Raw fastq reads for all ChIP-seq experiments were processed by using a snakemake-based pipeline (https://zenodo.org/record/819971). The raw reads were first processed by using FastQC(50), and uniquely mapped reads were aligned to the hg19 reference genome by using Bowtie(51) Ver. 1.1.2. Duplicate reads were removed by using SAMBLASTER(52) before compression to bam files. To directly compare ChIP-seq samples, uniquely mapped reads for each mark were downsampled per condition to 15 million, sorted, and indexed with the use of samtools Ver. 1.5(53). To visualize ChIP-seq libraries on the IGV genome browser, we used deepTools Version 2a.4.0(54), which generated bigWig files by scaling the bam files to reads per kilobase per million (RPKM).

Super ChIP-seq tracks were generated by merging bam files from each phenotype, sorting and indexing by using samtools, and scaling to RPKM by using deepTools. We used the Model-based analysis of ChIP-seq (MACS) version 1.4.2(55) peak calling algorithm to identify peaks for the sharp marks H3K4me1, H3K27ac, and H3K4me3; we used MACS version 2.1.0 to identify peaks for broad marks H3K9me3, H3K79me2, and H3K27me3. The peak enrichment was calculated over a whole genome “input” background with a p-value of < 1e-5. Super-enhancers were identified by using the ranking of the super-enhancers (ROSE) algorithm based on the H3K27ac ChIP-seq data(16).

### ChromHMM analysis

ChromHMM(14) is used to identify combinatorial chromatin state patterns based on the histone modifications studied. Normalized bam files were converted to bed files and binarized at a 1000-bp resolution by using the BinarizeBed command. We programmed ChromHMM to Learn 10 (**Fig. 1A**), 12 (**Fig. S1A**, left panel), and 15 (**Fig. S1A**, right panel) chromatin state models and chose the 10-state model for in-depth characterization and presentation purposes. We chose this model because a lot of the states were replicated in the higher state models, and the 10-state model was large enough to identify important functional elements yet small enough to allow easy interpretation of the biological states. Any marks with an emission probability of <0.1 (or 10%) were not considered for the annotation of the states.

To determine chromatin state differences between various groups, we used a two-step process using ChromXploreR (**Fig. S1C**). First, using the segmentation calls from the ChromHMM output, the entire genome was divided into non-overlapping bins of 1000 bp. We next counted the number of times a chromatin state was observed in each of the bins and obtained a frequency matrix for each state across all of the samples in the ChromHMM model (E1-E10). For visualization purposes, the frequency matrix was binarized such that if a bin contained a non-zero value for a sample, it was counted as 1, otherwise as 0. The top 10,000 most variable bins were used for further analysis. Most variable bins or regions were computed with the use of the *FSbyVar* function from the CancerSubtype package(56) in R. The distance matrix was calculated by using the Jaccard distance metric. The following distance matrix was used to generate Multidimensional scaling (MDS) plots to visualize the differences between the groups of samples (Fig 1B and Fig S2D-S2E). In the second step, low variable genomic loci were removed from the frequency matrix, and significant differences between two groups of sample types were calculated by using a nonparametric Mann Whitney Wilcoxon test with a p-value < 0.05 for each state separately (Fig S2F).

### DiffBind Analysis

The DiffBind R package(41) was used to find differences between two groups of samples based on singular marks. For each histone modification mark, the consensus peaks across all the tumors samples were obtained using the *dba* and *dba*.*count* function in the package with default options except for summit size, which was adjusted to 500. The matrix of the normalized reads was obtained across all the samples. Most variable bins or regions were computed with the use of the *FSbyMAD* function from the CancerSubtype(56), and the top 10,000 most variable loci were selected for further analysis. MDS plots were generated to visualize the differences between the tumor and normal samples (Fig S2A-S2B).

Similarly, a matrix of the top variable regions for H3K27AC mark (typical enhancers) was constructed across all the tumor (n = 33) and adenoma samples (n = 4) for consensus NMF (non-negative matrix factorization) analysis. Differentially expressed regions were calculated between tumor and normal samples using *dba*.*contrast* (default settings) and *dba*.*analyze* (arguments: method=DBS_DESEQ2, bSubControl=T) function in the DiffBind package (Figure 2A).

### NMF clustering into Epigenomic subtypes (EpiC)

Consensus NMF analysis was performed on the matrix of enhancers obtained from the DiffBind analysis using *nmf* function (arguments -- rank = 2:10, nrun=1000 and method=brunet (57) from the NMF(58) package in R. Based on the dispersion, cophenetic and silhouette scores, *k*, the optimal cluster number of four was determined (Fig S7A, Fig 3A).

### RNA-Access Sequencing and analysis of MDACC tumor and adjacent normal samples

mRNA libraries of the tumor (n = 7) and normal (n = 5) samples were prepared from 200 ng of total RNA with the use of the TruSeq Stranded mRNA HT sample preparation kit. Samples were dual-indexed before pooling. Libraries were quantified by qPCR with the use of the NGS Library Quantification Kit. Pooled libraries were sequenced by using the HiSeq2000 (Illumina) according to the manufacturer’s instructions. An average of approximately 30 million paired-end reads per sample were obtained. The quality of raw reads was assessed by using FastQC (https://www.bioinformatics.babraham.ac.uk/projects/fastqc/). The average per-base quality score across all of the files was greater than 32 and had passed all major tests. The raw reads were aligned to the *Homo sapiens* genome (hg19) using STAR v2.4.2a(59) (https://github.com/alexdobin/STAR/releases/tag/STAR_2.4.2a). The mappability of unique reads on average was ∼82% RNA-seq dataset. The raw counts were computed with the use of quantMode function in STAR. The read counts that were obtained are analogous to the expression level of each gene across all the samples. The differential expression analysis was done by using DESeq2(60). Genes with raw mean reads of greater than 10 were used for normalization and differential gene expression analysis using DESeq2 package in R. A Wald test, defined in the DESeq function of the package, was used for differential expression analysis, and shrunken log fold-changes (i.e., obtaining reliable variance estimates by pooling information across all the genes) were calculated. Genes with absolute log2 fold-change greater than log2(1.5) and padj < 0.05 were called as differentially expressed genes. Principal Component Analysis (PCA), MDS, hierarchical clustering plots (hclust), and XY plots between replicates of the same group or phenotype were used to examine for nominal amounts of nontechnical variation and other latent factors. PCA, hclust, and volcano plots were generated by using ggplot2 (Wickham). All of the heatmaps and box plots of differentially expressed genes were generated with the use of the pheatmap package (https://cran.r-project.org/web/packages/pheatmap/pheatmap.pdf) and ggpubr package (https://rpkgs.datanovia.com/ggpubr/), respectively.

### RNA-seq analyses of TCGA COAD and Adenoma dataset

The COAD TCGA(24) RNA-seq version 2 level 3 normalized dataset (COAD.rnaseqv2 illuminahiseq_rnaseqv2 unc_edu Level_3 RSEM_genes_normalized_ _data.data) was downloaded from http://gdac.broadinstitute.org/. Only primary tumors (n = 285) and normal samples (n = 41) were used for further analysis. The normalized matrix was used for differential gene expression analysis using DESeq2 package.

The transcriptome analysis of the polyp dataset(61,62) (GEO(63) accession numbers: GSE88945, GSE106500, and GSE136044) consisting of adenoma (n = 16) and normal crypts (n = 10) was performed similarly to RNA Access sequencing dataset. Briefly, the dataset was mapped to the hg19 genome using STAR aligner (v2.4.2a). The mappability of unique reads on average was ∼90% RNA-seq dataset. Differential expression analysis was performed using the DESeq2 package. Genes with absolute log2 fold-change greater than log2(1.5) and padj < 0.05 were called as differentially expressed genes for both TCGA and adenoma dataset.

### HiChIP Analysis

HiChIP was performed as described(64). Briefly,1 ⨯ 10^7^ SW480 cells were crosslinked. In situ contacts were generated in isolated and pelleted nuclei by DNA digestion with MboI restriction enzyme, followed by biotinylation of digested DNA fragments with biotin–dATP, dCTP, dGTP, and dTTP. Thereafter, DNA was sheared with Covaris E220 with the following parameters: fill level = 10, duty cycle = 5, PIP = 140, cycles/burst = 200, and time = 4 min; chromatin immunoprecipitation was done for H3K27Ac with use of anti-H3K27ac antibody. After reverse-crosslinking, 150 ng of eluted DNA was taken for biotin capture with Streptavidin C1 beads followed by transposition with Tn5. In addition, transposed DNA was used for library preparation with Nextera Ad1_noMX, Nextera Ad2.X primers, and Phusion HF 2XPCR master mix.

The following PCR program was performed: 72 °C for 5 min, 98 °C for 1 min, then 11 cyclesat 98 °C for 15 s, 63 °C for 30 s, and 72 °C for 1 min. Afterward, libraries were two-sided size selected with AMPure XP beads. Finally, libraries were paired-end sequenced with reading lengths of 76 nucleotides. HiChIP paired-end reads were aligned to the hg19 genome; duplicate reads were removed, assigned to MboI restriction fragments, and filtered for valid interactions. Interaction matrices were generated with *HiC-Pro*(65). HiC-Pro filtered reads were then processed, and loops were identified with the use of *hichipper*(66).

### 3C-qPCR Validation

Chromosome conformation capture (3C) was performed as previously described by Hagège et al. (67) and Fullwood et al. (68) for SW480 cells. Briefly, 1 ⨯ 10^7^ cells were crosslinked with 1% formaldehyde and quenched with 125 mM glycine; the obtained nuclei and a bacterial artificial chromosome (BAC) clone were subsequently digested with 400 U of DraIII, followed by ligation, reverse-crosslinking, and DNA purification. DraIII, a 6-base cutter, was selected because it allowed for individual analysis of the designated enhancers (ASCL2 E1-E4) and was deemed appropriate for the large locus size. The BAC DNA (source: CHORI) spanned the entire locus of interest and served as a control template.

After the generation of the 3C libraries, RT-qPCR was used to determine the relative interaction frequencies. 3C primers were designed according to previous guidelines(69). Genomic coordinates for the active enhancers of ASCL2 were filtered by H3K27ac peaks; test primers for the four predicted enhancers were selectively designed based on this information. A random test primer was designed at a DraIII site upstream of E1 (designated T1). Test primers were coupled with an anchor primer that represented the ASCL2 TSS to assess enhancer-promoter interactions. A gene desert (i.e., “gene poor”) within chromosome 11 was used as an internal control to normalize 3C-qPCR data. Normalized relative interaction was calculated as 2^-ΔCt^= [(C_t interaction_ – C_t gene desert_)_3C_ – (C_t interaction_ – C_t gene desert_)_BAC_].

### Motif analysis and TF enrichment

We performed de novo motif discovery and a known matching motif with the use of the HOMER(25) pipeline (http://homer.ucsd.edu/homer/motif/) using default settings and background file that consisted of all the peaks or genomic regions called in the analysis. For calling the enriched TFs for Figure 1I, the background file consisted of all the peaks called in CRC tumor and normal samples whereas, for Figure 3D, the background file consisted of the enriched differentially expressed regions across all the four EPIC clusters.

### Peak length analysis

All peaks (called by MACS2 at p < 0.01) less that 1kb apart within a group (Epic3 and Epic4) were merged before generating a list of peaks that were present in 100% of samples within that group. For fig. 3D, we generated a kernel density estimate (KDE) of the average peak sizes (across samples) by group to assess peak length distribution. For fig. 3F,we filtered peaks based on average peak size across samples to generate bar plots of broad peaks (left, >5kb) and sharp peaks (right, <3kb)

### UpSetR

All the ChIP-seq peaks from the H3K27ac mark were associated with the nearest gene, and then the UpSetR(70) package was used to visualize intersections of these genes across the tumor and cell lines.

### Pathway Analysis

Enhancers were assigned to the genes using GREAT(17) and previously published enhancer-target networks inferred by JEME from the ENCODE+Roadmap and FANTOM5 dataset(18). Pathway analysis was performed using the Consensus Molecular Database Pathway analysis tool(71).

### CMS2 Classification into EpiC3 and EpiC4

115 tumor profiles were identified as CMS2-epithelial/canonical subtype on Illumina HiSeq RNA-Seq data from The Cancer Genome Atlas (TCGA). EpiC 3 and EpiC 4 genes were determined from enhancer-gene annotations as described above under ‘pathway analysis’. We downloaded raw gene expression data and apply across-sample normalization based on variance-stabilizing transformation (VST) (60). Three molecular subtypes were validated based on cophenetic scores in 115 cancer cases using 2098 EpiC 3 / EpiC 4 gene signatures for consensus non-negative matrix factorization (NMF) clustering. Clinical data for all profiles were downloaded from the TCGA data portal and the prognosis of each cluster was examined by Kaplan-Meier overall survival estimators. The survival outcomes of each cluster were compared by log-rank tests (survminer R package). Survival analysis between cluster 2 and cluster 3 (100 total samples out of 115 samples) revealed reduced overall survival for cluster 3.

### Enhancer modulation using CRISPR-dCas9-KRAB

In order to modulate gene expression without altering the target DNA sequences, an RNA-guided, catalytically inactive Cas9 (dCas9) fused to a transcriptional repressor domain (KRAB) was used to silence genomic regions identified as enhancers via KRAB repression at the promoter region. To generate a dCas9-KRAB effector stable cell line, we produced lentiviral particles from pHAGE EF1α dCas9-KRAB (Addgene plasmid # 50919) or pHAGE TRE dCas9-KRAB (Addgene plasmid # 50917) using a standard protocol. Transduced cells were selected for 6 days with the use of antibiotic resistance and were expanded to generate a stable cell line.

Next, gRNAs were designed by using the GPP Web Portal of the Broad Institute. Annealed gRNA oligos were ligated to pLKO.1-puro U6 sgRNA BfuAI stuffer (Addgene plasmid # 50920) or to pLenti SpBsmBI sgRNA Hygro (Addgene plasmid # 62205), and lentiviral particles were generated. A transduction procedure was performed in the stable dCas9-KRAB cell line, and transduced cells having both dCas9-KRAB and gRNA constructs were selected with the use of antibiotic resistance.

To evaluate the effects of the recruitment of dCas9-KRAB to the target enhancer’s genomic region, H3K27ac or H3K4me1 ChIP followed by quantitative PCR for enhancer regions was performed to assess the enrichment level of H3K27ac or H3K4me1 at the enhancer site in modulated cells compared with the non-modulated parental control cells. To investigate the impact of enhancers’ modulation on the corresponding gene expression, qRT-PCR was performed for the target gene.

### Proliferation and invasion assays

Cells were seeded in 180 µl of medium in 96-well plates. Plates were maintained at standard culture conditions at 37 °C without CO_2_ and then 20 µl of alamarBlue (Sigma) was added to the cells. The magnitude of the reduction of alamarBlue correlates with the number of viable cells. Fluorescence of each sample was measured at an excitation wavelength of 560 nm and emission of 590 nm with a Tecan Infinite 200 spectrophotometer at various time points (0, 24, 48, 72, 96, 120 h).

Cell invasion was assessed by using a Tumor Invasion System, 24-Multiwell Insert Plate (BD Biosciences). A BD Falcon FluoroBlok 24-Multiwell Insert System was used as a control for cell migration. This assay was performed according to the manufacturer’s protocol. Fluorescence was measured after 24 h by using a Tecan Infinite 200 microplate reader at an excitation/emission of 494 nm/517 nm. Experiments were performed in triplicate.

### *In vitro* small molecule inhibitor assays and combination therapies

Small molecule inhibitors, as shown in various experiments in Figure 4, and supplementary figures S10, S11, S12, S13 and S14 were used in CRC cell lines. Inhibitors were used in final concentrations of 100 pM, 1 nM, 10 nM, 100 nM, 1 μM, and 10 uM. A cell viability assay was performed as described previously(72,73). IC_50_ and AUC were calculated for each compound in each of the cell lines. Based on these indices, the most potent/efficient compounds were used for combinatorial treatments.

### Mouse experiments

#### Patient-derived xenografts

Primary human-tumor xenograft models were established as previously described(74). Tumor specimens were obtained from patients with metastatic colorectal cancer being treated at The University of Texas MD Anderson Cancer Center, and all patients provided informed written consent for specimens to be used for research purposes including implantation in xenografts. Samples were obtained with the approval of the Institutional Review Board. Xenografts were established in NOD/SCID 8-week-old female mice. Once established, PDXs were expanded in 69 athymic nude mice for experiments.

After tumors were established with the median tumor volume exceeding 300 mm^3^, treatment of 3-6 mice per arm was commenced via oral gavage with either vehicle control (0.5% Tween 80/0.5% CMC in water), binimetinib (3 mpk, BID), trametinib (0.25 mpk, QD), iBET-151 (15 mpk, QD), or drug combinations as indicated in Figures 2I-2J. Tumor size and mouse weight were measured twice a week. After 21 days, treatment was discontinued, and mice were euthanized. Tumors from 3 mice per arm were excised (2 - 4 hours after treatment), segmented, and immediately flash-frozen in liquid nitrogen or in 10% buffered formalin solution.

#### Cell line xenografts

HCT116 (CMS1), T84 (CMS2), LS513 (CMS3), and SM620 (CMS4) were grown in the manufacturer-recommended media. One million cells were transplanted per site, and 5 mice were injected per experimental arm. Once tumors reached ∼5mm in size, the treatment was started as indicated in the specific figures/legends. All drugs were purchased from commercial sources: i-BET151 (MedChemExpress, 15mg/kg, every other day), gefitinib (EGFRi, Sigma, 100mg/kg, every other day), SB431542 (TGFβi, MedChemExpress, 10mg/kg, every other day), and olaparib (APExBio, 50mg/kg, every other day) and dissolved in the manufacturer-recommended vehicles. 30% PEG-300 was used as the vehicle control. Tumor volume was measured every other day. Once tumors reached 1.5 cm in one direction in any of the experimental arms, the mice were euthanized and the tumors were harvested.

### Production of anti-LGR5 mAb

Cloning and production of anti-LGR5 (8F2) mAb rat/mouse IgG2a chimera was performed as previously reported (75).

### Generation of anti-LGR5 mAb-RVX-208 conjugate

**Scheme 1:**
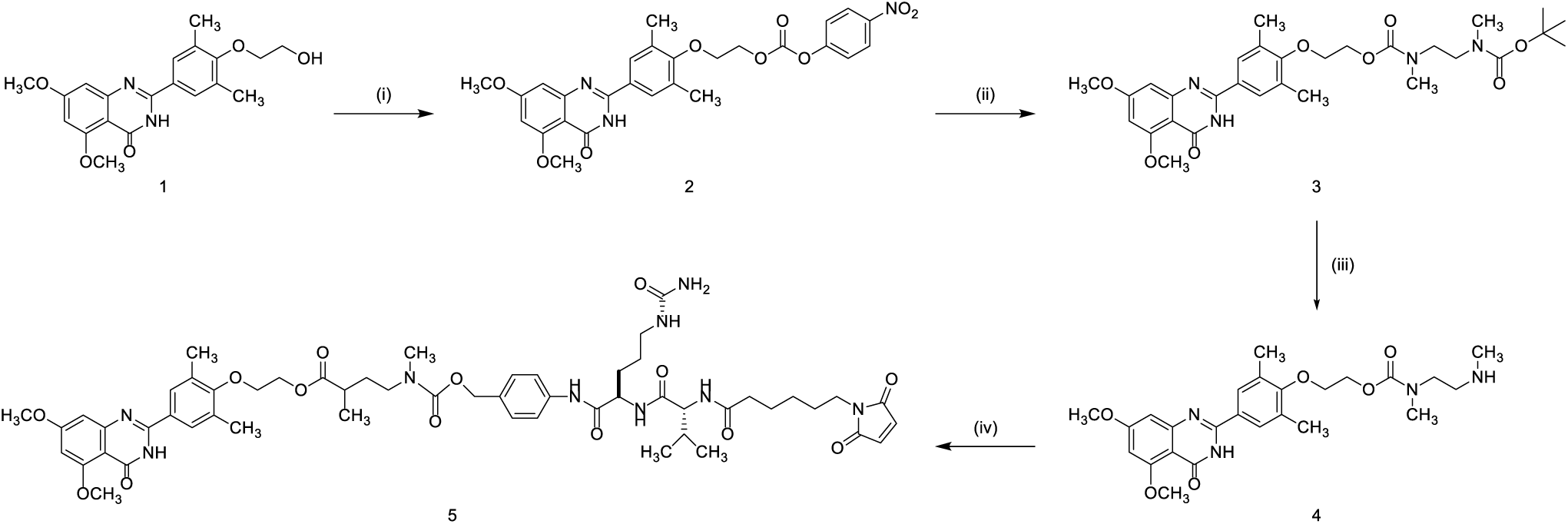
(i) Bis (4-nitrophenyl) carbonate, *i*-Pr_2_NEt, DMF, r.t., overnight stirring; (ii) *tert*-butyl methyl [2-(methylamino) ethyl]carbamate, 2,6-lutidine, 45°C, overnight stirring; (iii) TFA, CH_2_Cl_2_, r.t., 2 h stirring; (iv) MC-Val-Cit-PAB-PNP, *i*-Pr_2_NEt, 37°C, overnight stirring.

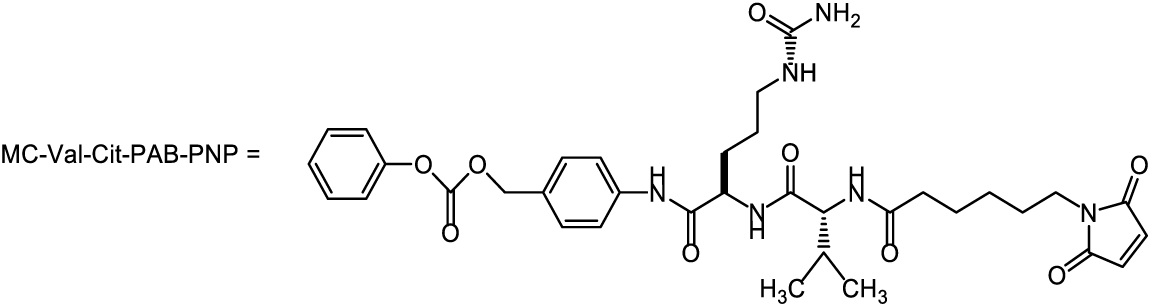

#### Synthesis of 2-(4-(5,7-dimethoxy-4-oxo-3,4-dihydroquinazolin-2-yl)-2,6-dimethylphenoxy)ethyl (4-nitrophenyl) carbonate (2)

A solution of RVX-208 (1) (20 mg, 0.054 mmol) and bis(4-nitrophenyl) carbonate (50 mg, 0.16 mmol) in anhydrous *N,N*-dimethylformamide (DMF) (3 mL) was mixed with 25 µL *N,N*-diisopropylethylamine (17 mg, 0.16 mmol) and stirred room temperature (r.t.) overnight. The reaction mixture was evaporated and purified by column chromatography (SiO_2_; 95% (vol/vol) CH_2_Cl_2_/MeOH) to give the expected product with high yield (23 mg, 70%). The product was verified by mass spectra analysis. MS, ESI+: *m/z* calculated for C_27_H_25_N_3_O_9_, 535.16; found *m/z*, 536.00 (M+H)^+^.

#### Synthesis of *tert*-butyl (2-(4-(5,7-dimethoxy-4-oxo-3,4-dihydroquinazolin-2-yl)-2,6-dimethylphenoxy)ethyl) ethane-1,2-diylbis(methylcarbamate) (3)

A solution of compound **2** (30 mg, 0.028 mmol) and *tert*-butylmethyl [2-(methyl amine) ethyl] carbonate (30 mg, 1.2 equivalent, 0.067 mmol) was prepared in anhydrous DMF (2 mL) and heated at 45°C. To the reaction mixture, 1-hydroxy-7-azabenzotriazole (HOAt) (10 mg, 0.073) and 2,6-lutidine (23 mg, 0.21 mmol) were added and heated overnight with continuous stirring. Following evaporation of DMF, the crude product was purified by column chromatography (SiO_2_; 98% (vol/vol) CH_2_Cl_2_/MeOH) to yield 28 mg of **3** (84% yield). MS, ESI+: *m/z* calculated for C_30_H_40_N_4_O_8_, 584.28; found *m/z*, 584.9 (M+H)^+^.

#### Synthesis of 2-(4-(5,7-dimethoxy-4-oxo-3,4-dihydroquinazolin-2-yl)-2,6-dimethylphenoxy)ethyl methyl(2-(methylamino)ethyl)carbamate (4)

Trifluroacetic acid (1 mL) was added to a solution of compound **3** (28 mg, 0.047 mmol) in anhydrous dichloromethane (2 mL) under N_2_ atmosphere and stirred at room temperature. After 2 h, the solvent was evaporated and the reaction mixture was washed with diethyl ether and dried to give compound **4** (21 mg, 90 % yield). MS, ESI+: *m/z* calculated for C_25_H_32_N_4_O_6_, 484.23; found *m/z*, 485.00(M+H)^+^.

#### Synthesis of RVX-208-MC-Val-Cit-PAB-PNP-maleimide (5)

To a 2 mL solution of anhydrous DMF containing MC-VC-PAB-PNP (30 mg, 0.0413 mmol) and compound **4** (20 mg, 0.0413 mmol), 30 µL of *N,N*-diisopropylethylamine (20 mg, 0.19 mmol) was added and the reaction was heated at 37°C overnight. The crude product was purified on HPLC and to give 24 mg of **5** (55 % yield). MS (ES) m/z: 1082.8[M+H]^+^; 1104.9[M+Na]^+^ MS, ESI+: *m/z* calculated for C_54_H_70_N_10_O_14_, 1082.51; found *m/z*, 1082.8 (M+H)^+^,1104.9 (M+Na)^+^.

**Scheme 2:**
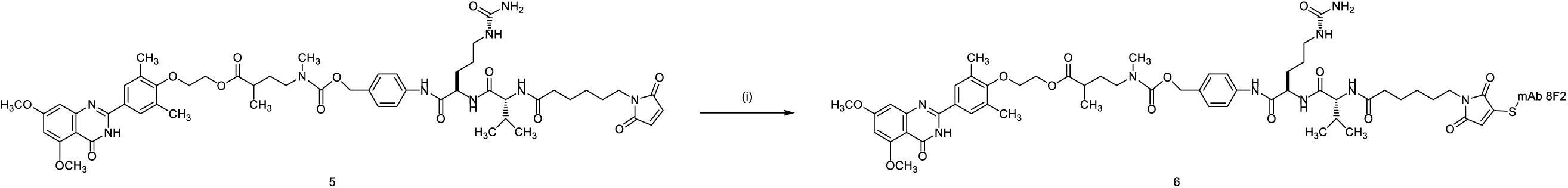
(i) Reduced 8F2, DMSO, Na_2_PO_4_ buffer (pH 8.2), 3 h at r.t., then 4°overnight.

#### Synthesis of anti-LGR5 (8F2) antibody-drug conjugate (6)

To a solution of 8F2-mAb (270 µg, 0.0018 µmol) in PBS (170 µL), 5 molar equivalent of DTT (30 µg, 0.19 µmol, 30 µL H_2_O) was added to partially cleave the interchain disulfides. The reaction was warmed to 37°C and proceeded for 3 h. After cooling to r.t., the mixture was purified with a Zeba desalting spin column in Na_2_PO_4_ (pH 8.2) buffer at a concentration of 1 mg/mL. A 10 molar equivalent of 5 (20 µg, 30 µL) in DMSO was added to the mAb solution and continuously stirred for 3 h at r.t., and then at 4°C overnight. The resulting ADC, **6**, was purified with a Zeba desalting spin column and collected in PBS at a final concentration of (1 mg/mL).

## Code availability

The ChIP-seq analysis source code are available at the following webpage: https://doi.org/10.5281/zenodo.819971.

## ACKNOWLEDGEMENTS

We thank Integromics group, the Advanced Technology Genomics Core Facility (NCI Grant CA016672(ATGC), the Research Animal Support Facility, the Advanced Microscopy Core Facility (funded by NIH S10 RR029552), Functional Genomics Core (NCI Cancer Center Support Grant (P30 CA016672) and MD Anderson Cancer Center Clinical Core. This work was supported by ACS research scholar award, CPRIT IIRA award (RP200390), a Career Development Award from MDACC GI SPORE to K.R. We thank Veena Kochat, Sharon Landers and Angela Bhalla for helpful discussions and suggestions.

## AUTHOR CONTRIBUTIONS

EO conceived, conceptualized and designed the study, planned and carried out experiments, analyzed data, made figures, and wrote the manuscript. ATR conceptualized and designed the study, carried out computational analysis, prepared figures, and wrote the manuscript. AKS carried out experiments, computational analysis and made figures regarding HiChIP and wrote the manuscript. A.S. performed mouse treatment experiments. EA performed computational analysis for CMS2 classification. AKG performed the 3C experiments and edited the manuscript. JS performed peak length analysis. MT helped with informatics analysis. CJT, SCC, MM, KT, ZJ, JSD, SG, HML, LRU, KC, YL, HC, AA provided technical help. JSM, EV, KSC and SK contributed to study design and provided reagents. KR conceived, conceptualized and designed the study, performed experiments, evaluated data, made figures and wrote the manuscript. All the authors edited the manuscript.

## FIGURE LEGENDS

**Figure S1.**
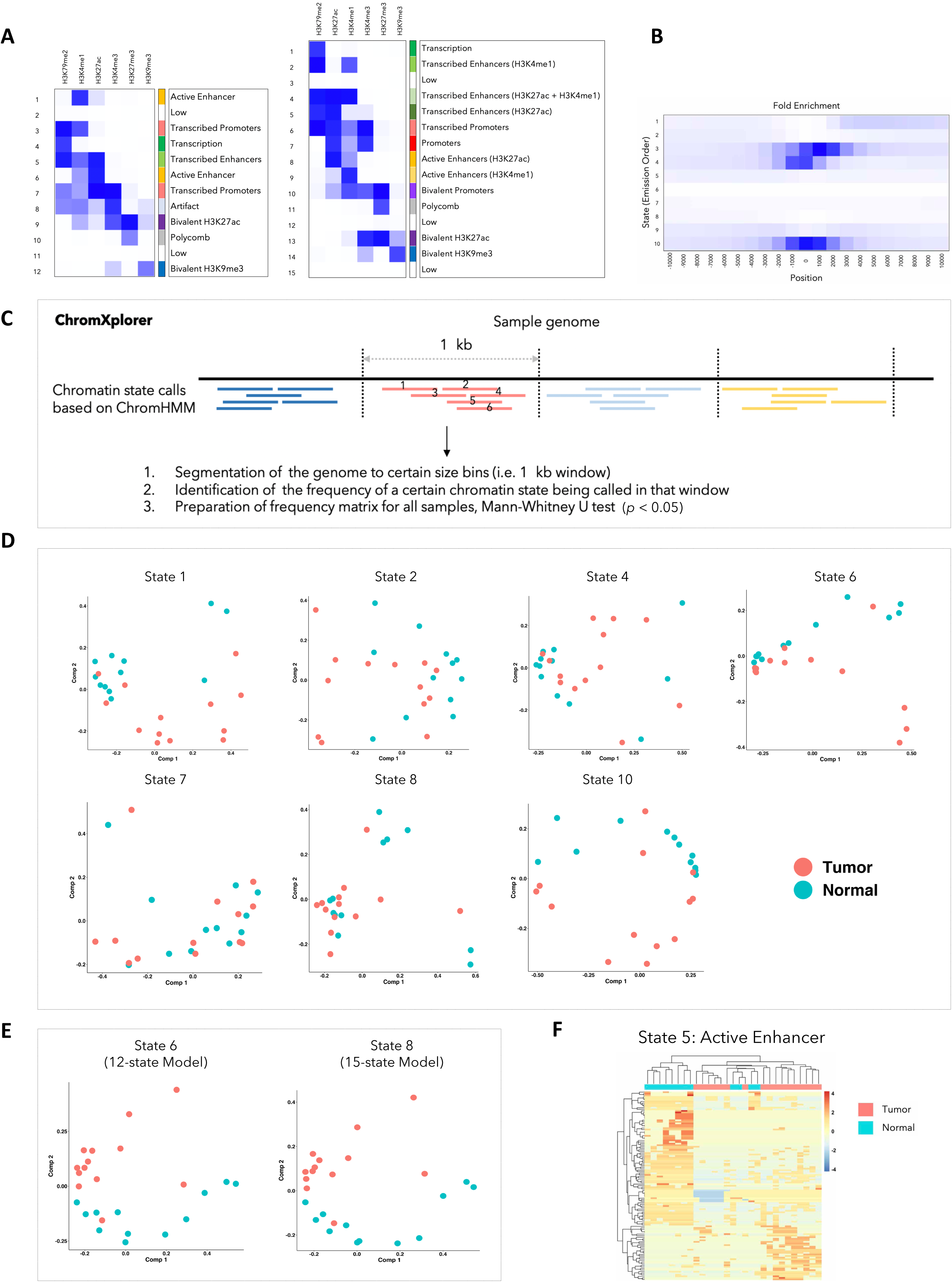

**Figure S2.**
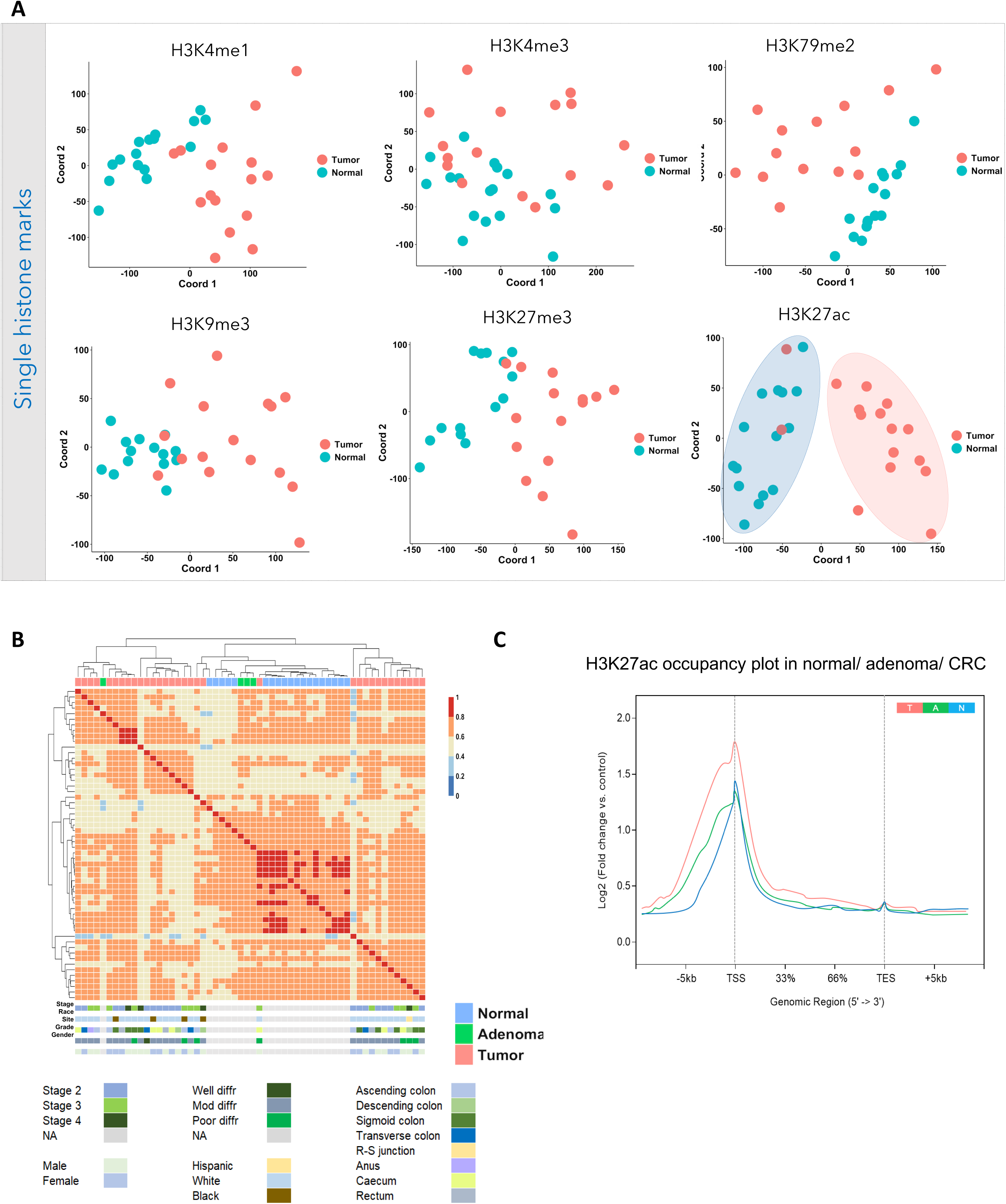

**Figure S3.**
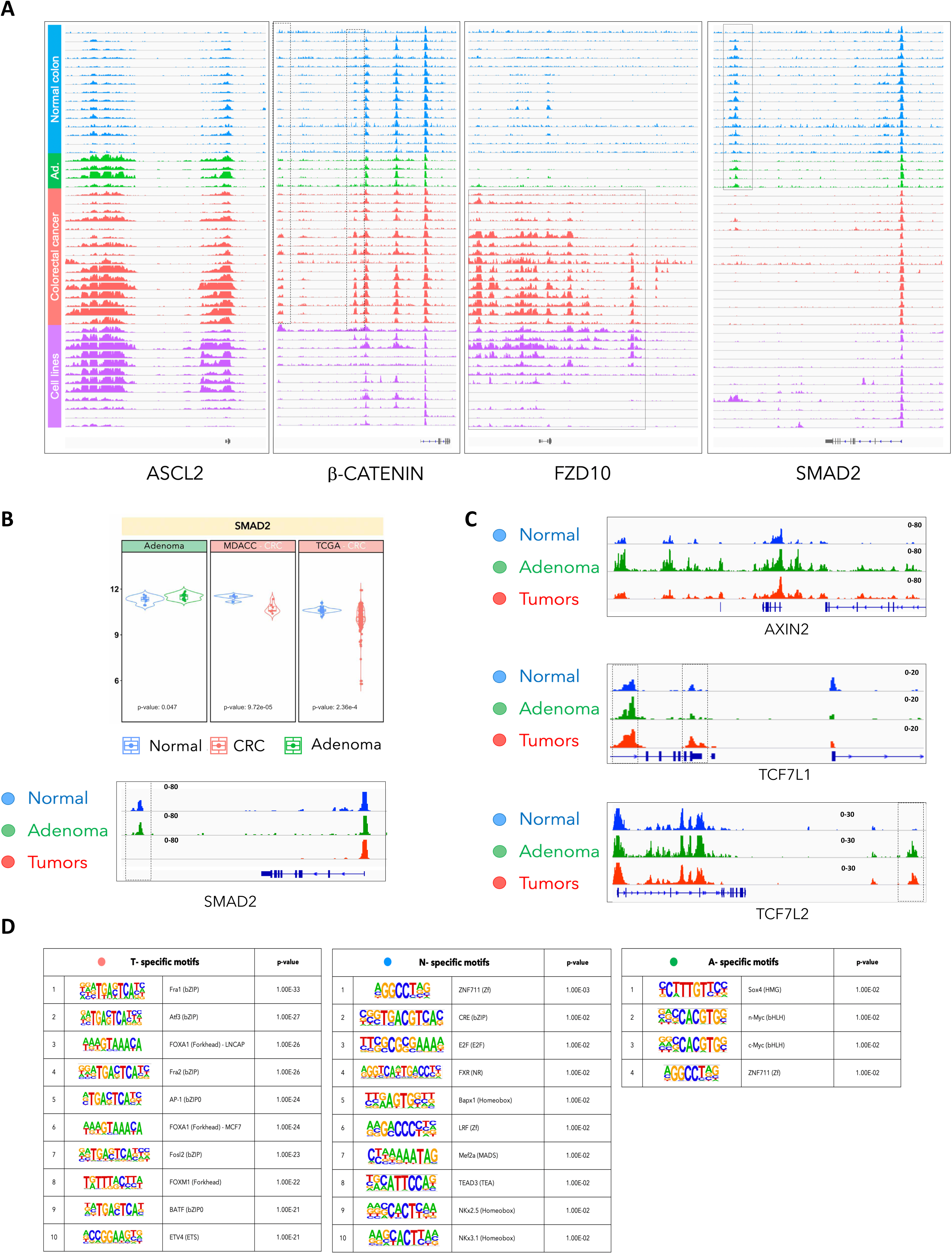

**Figure S4.**
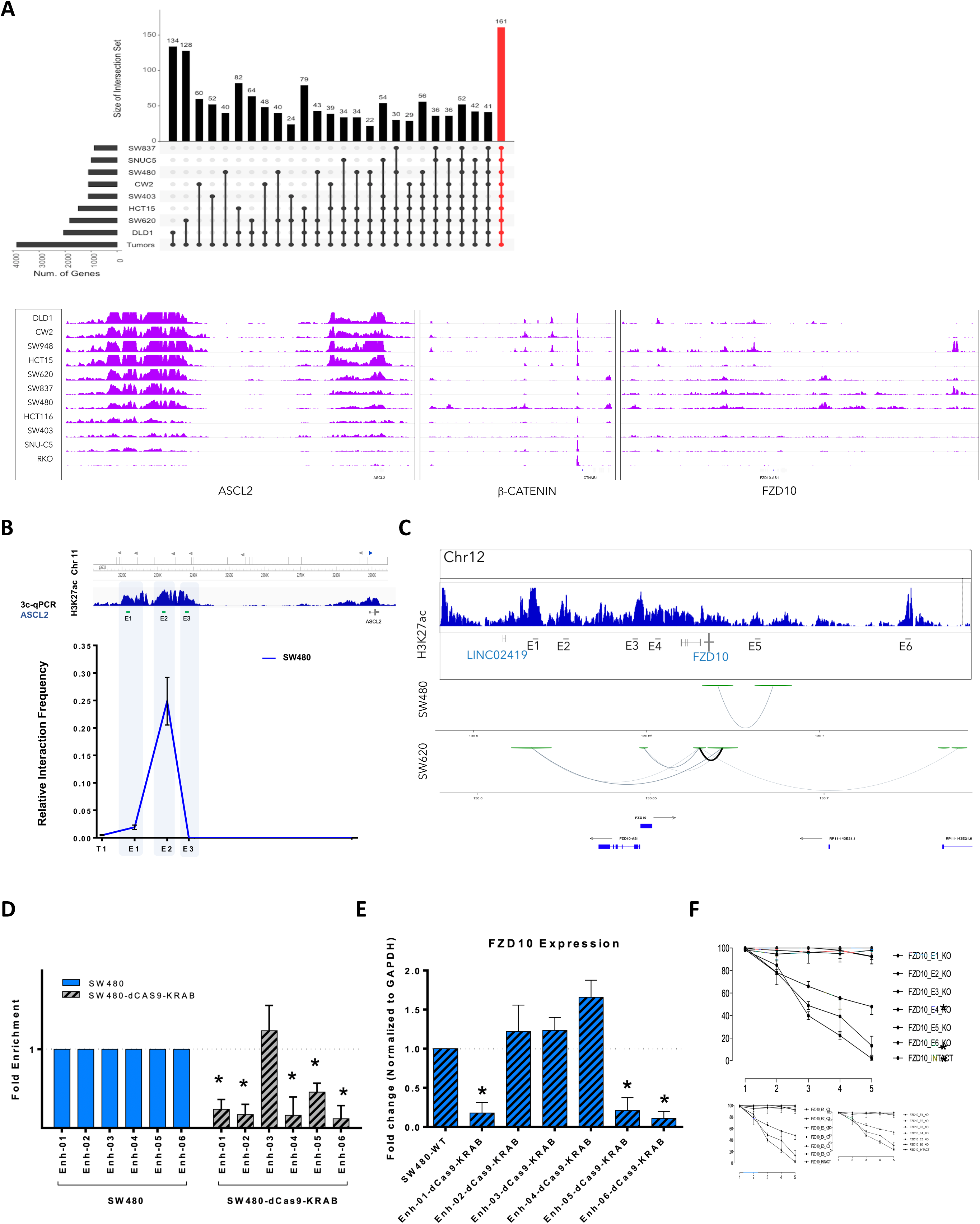

**Figure S5.**
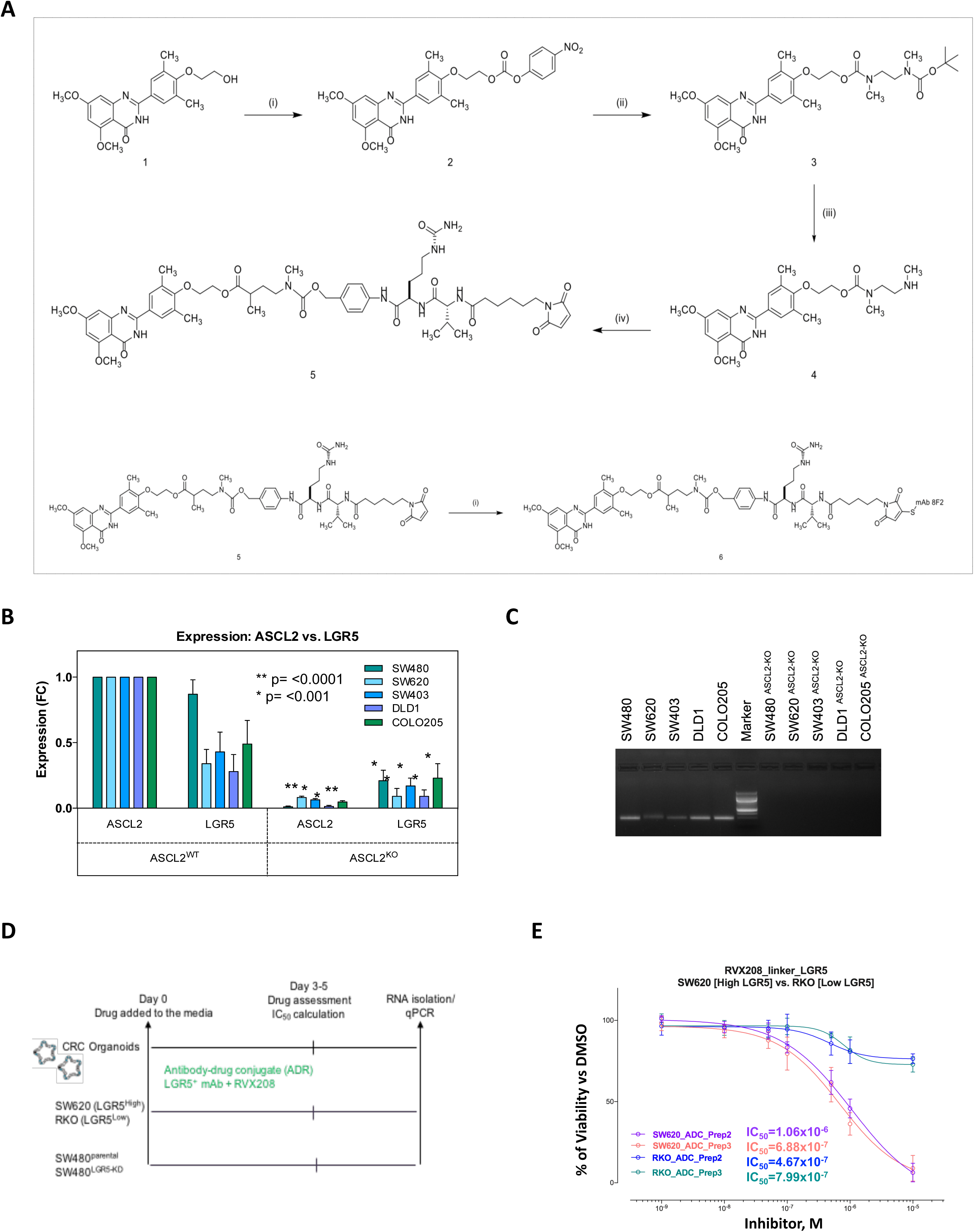

**Figure S6.**
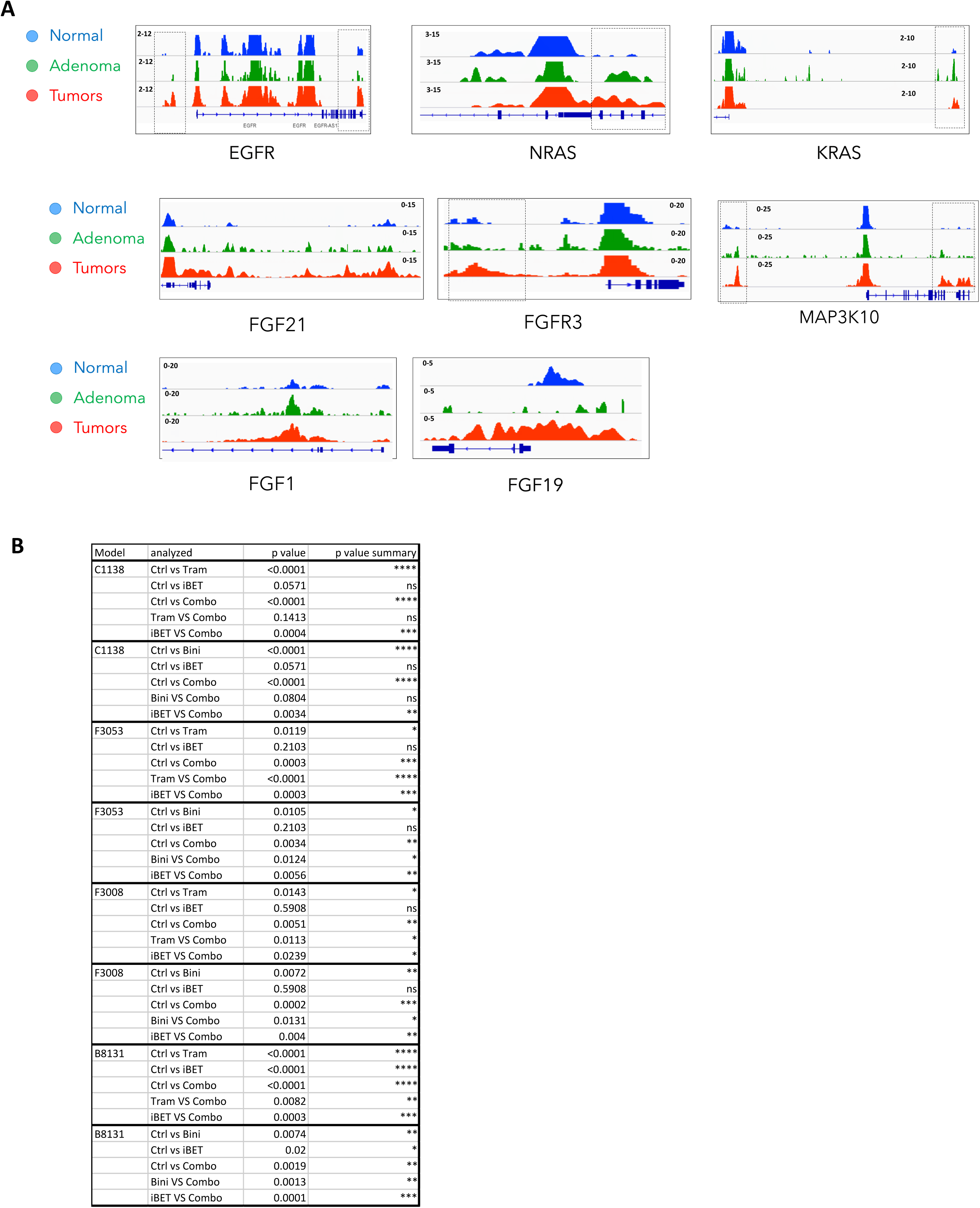

**Figure S7.**
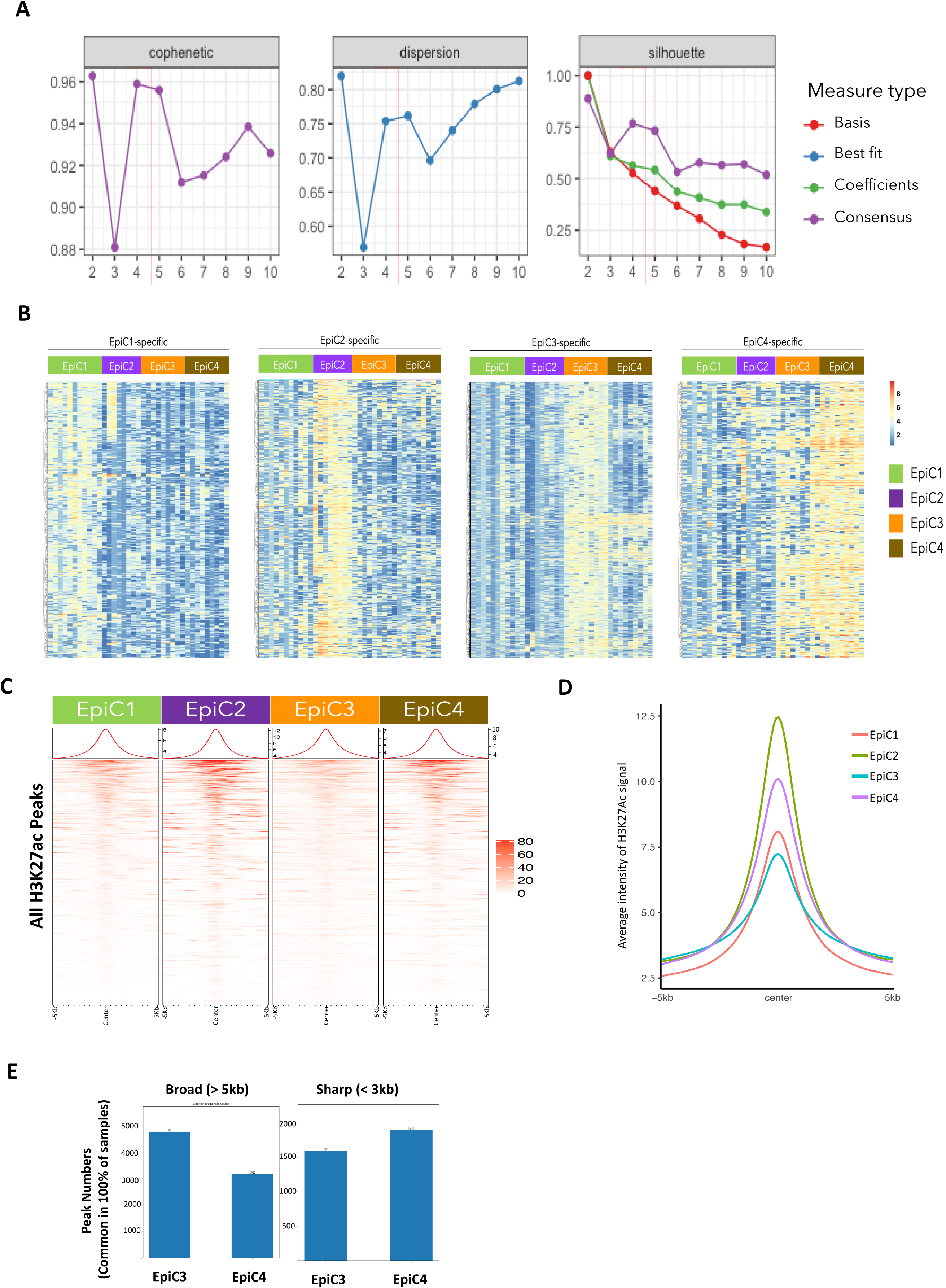

**Figure S8.**
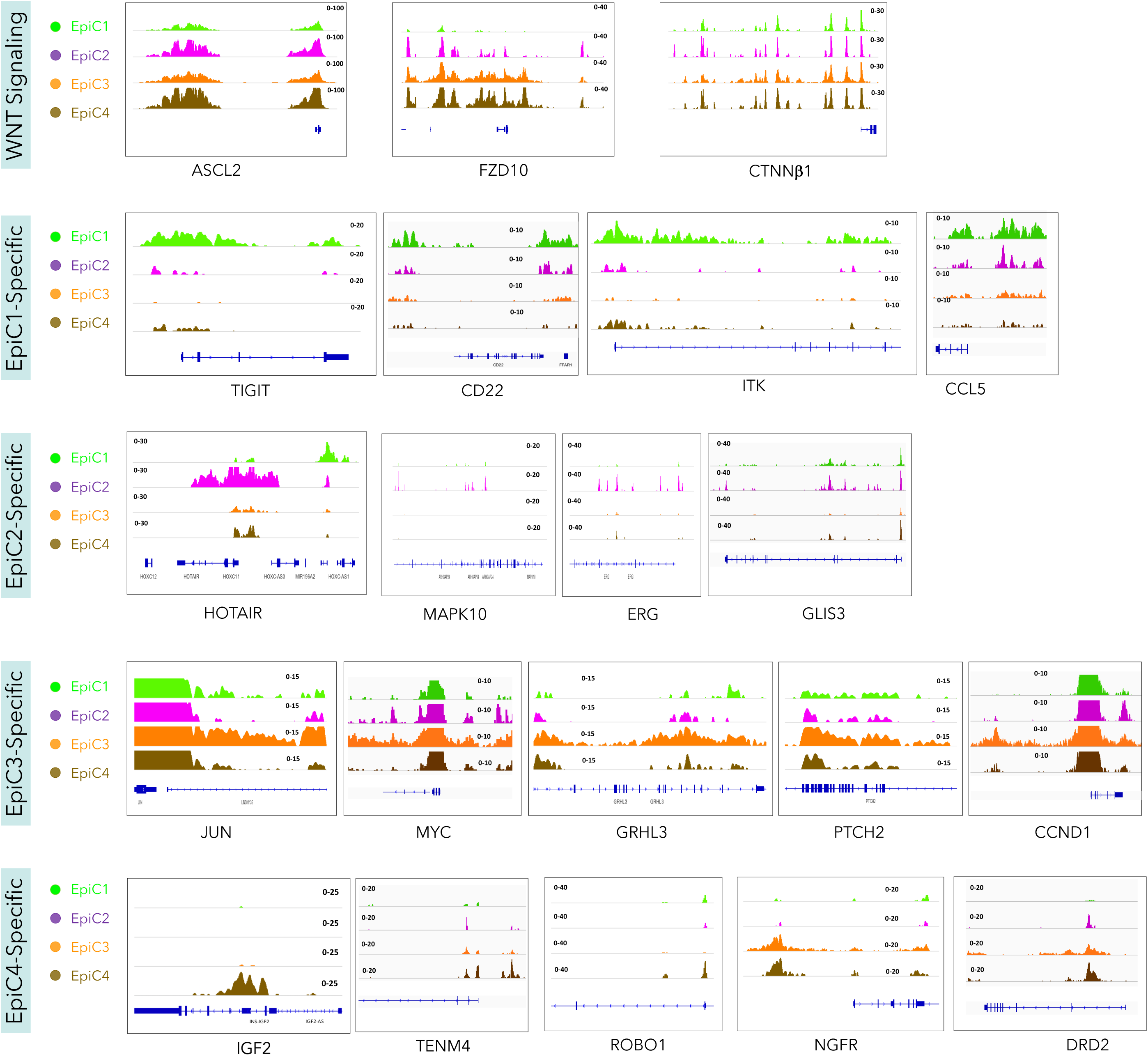

**Figure S9.**
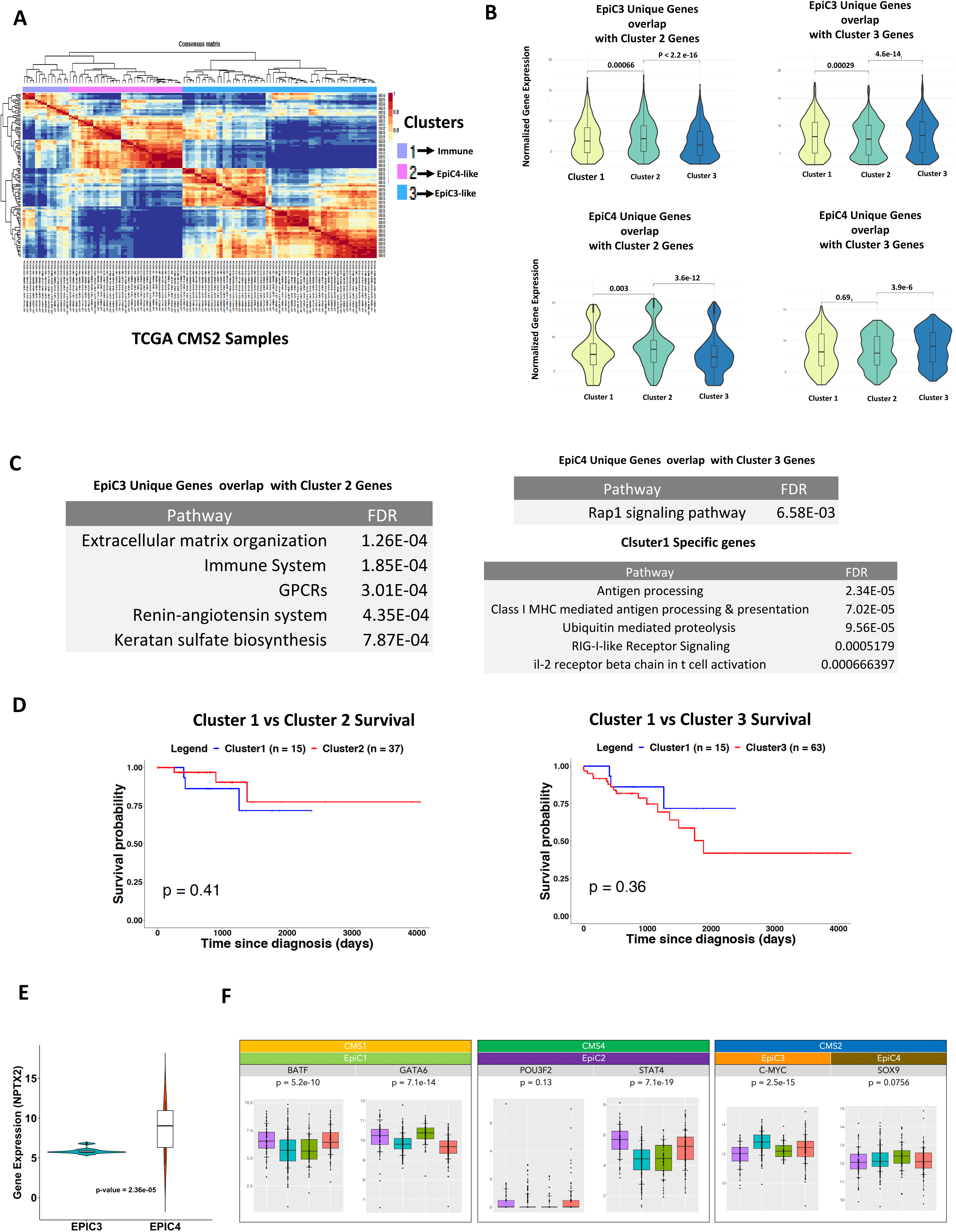

**Figure S10.**
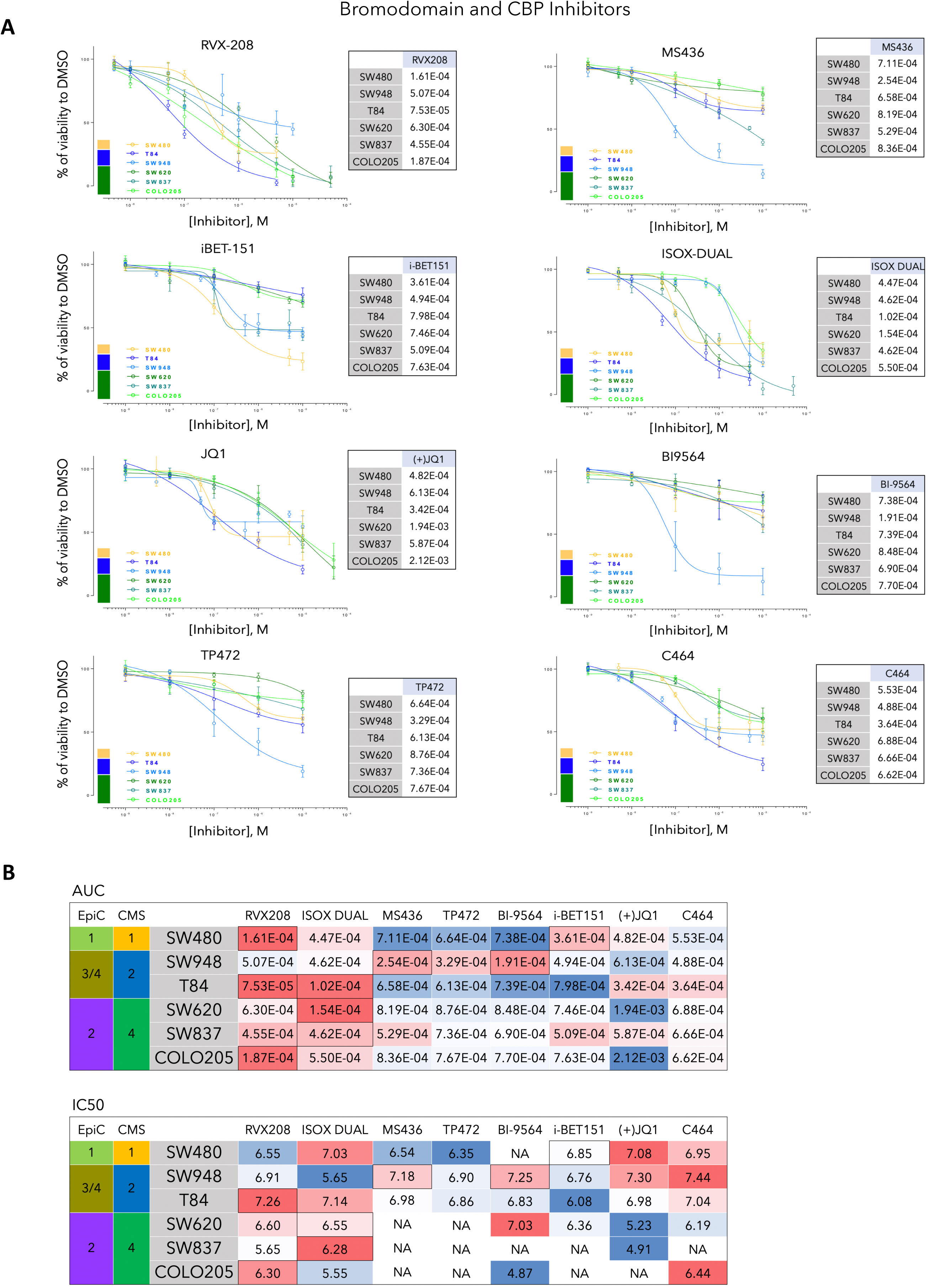

**Figure S11.**
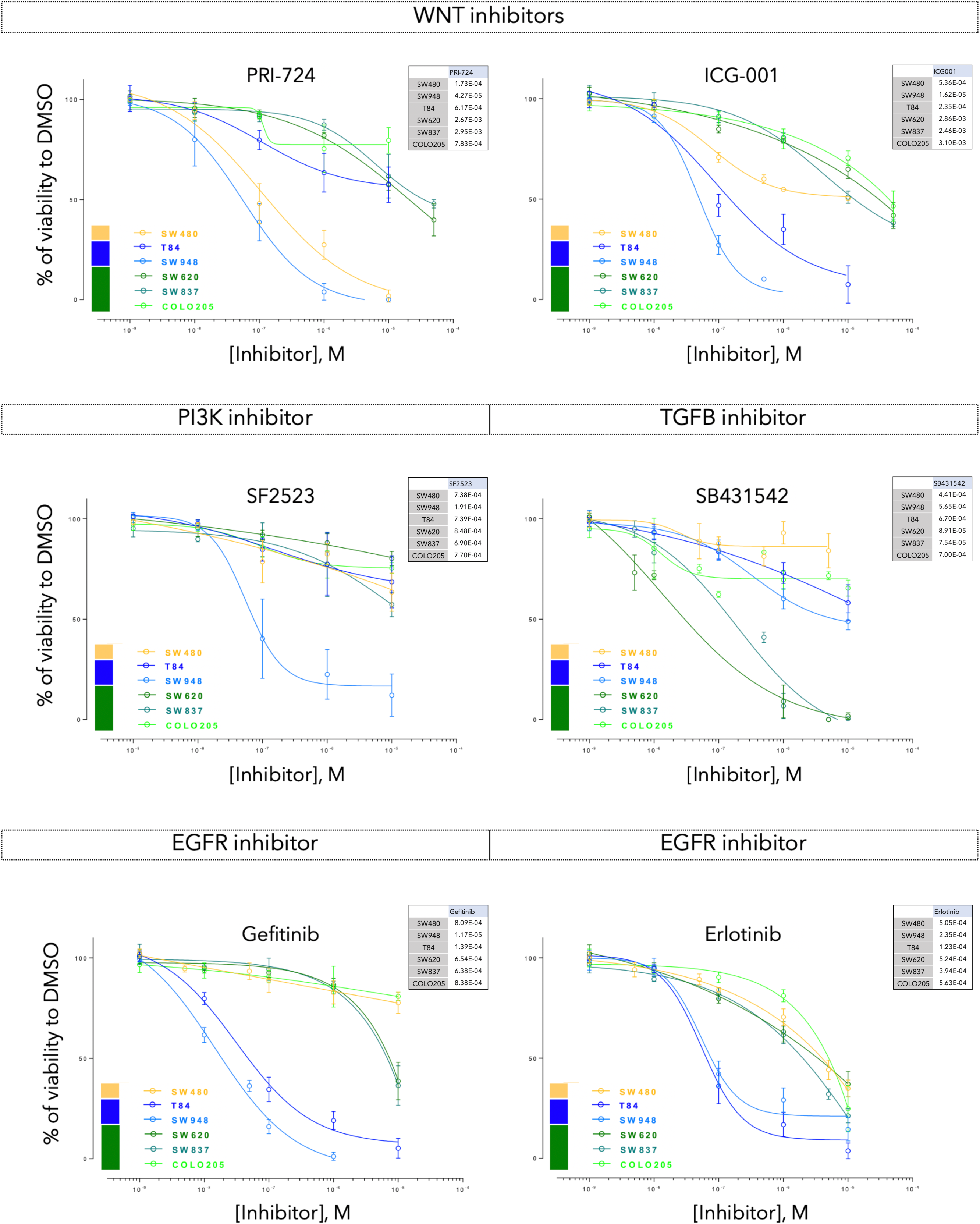

**Figure S12.**
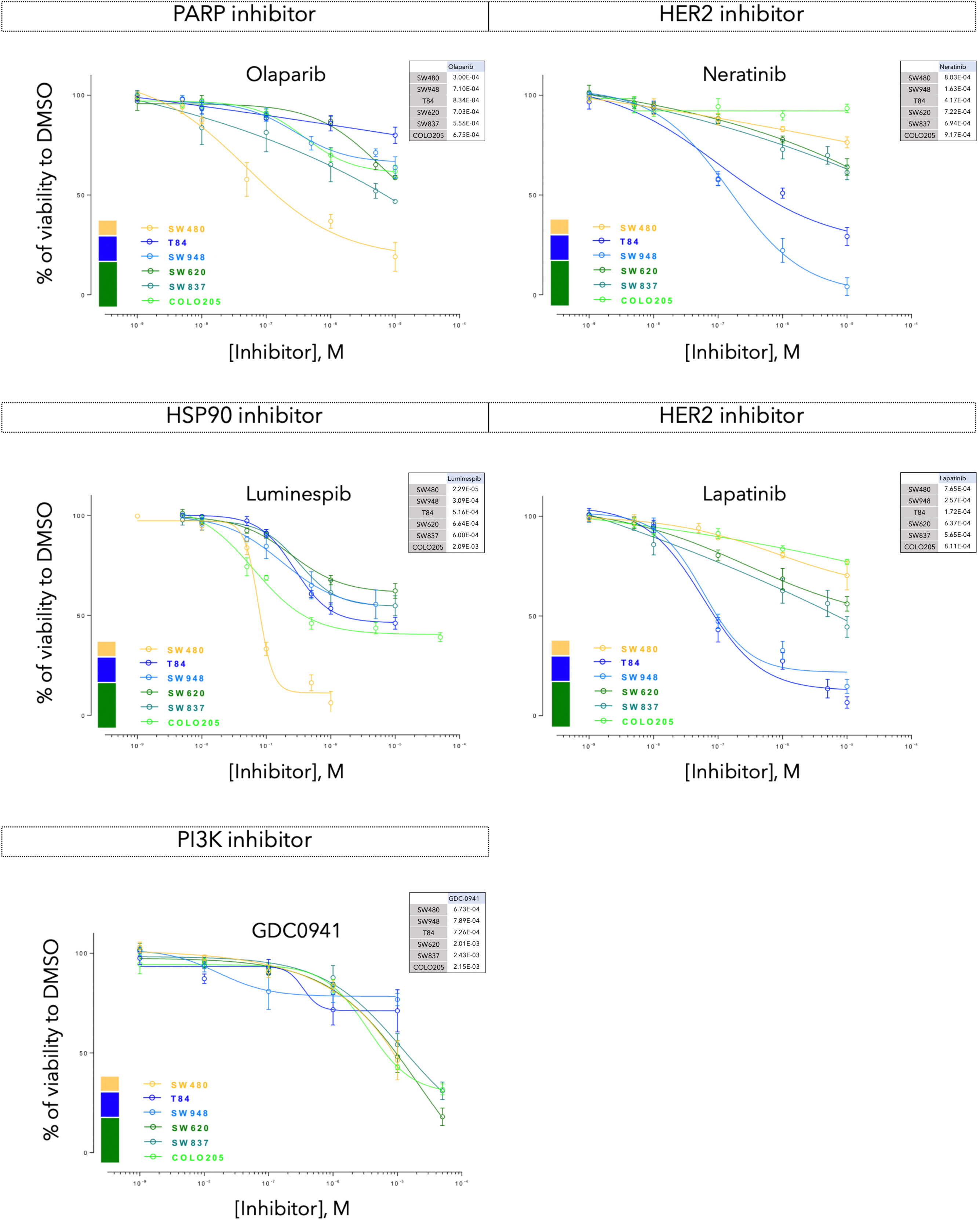

**Figure S13.**
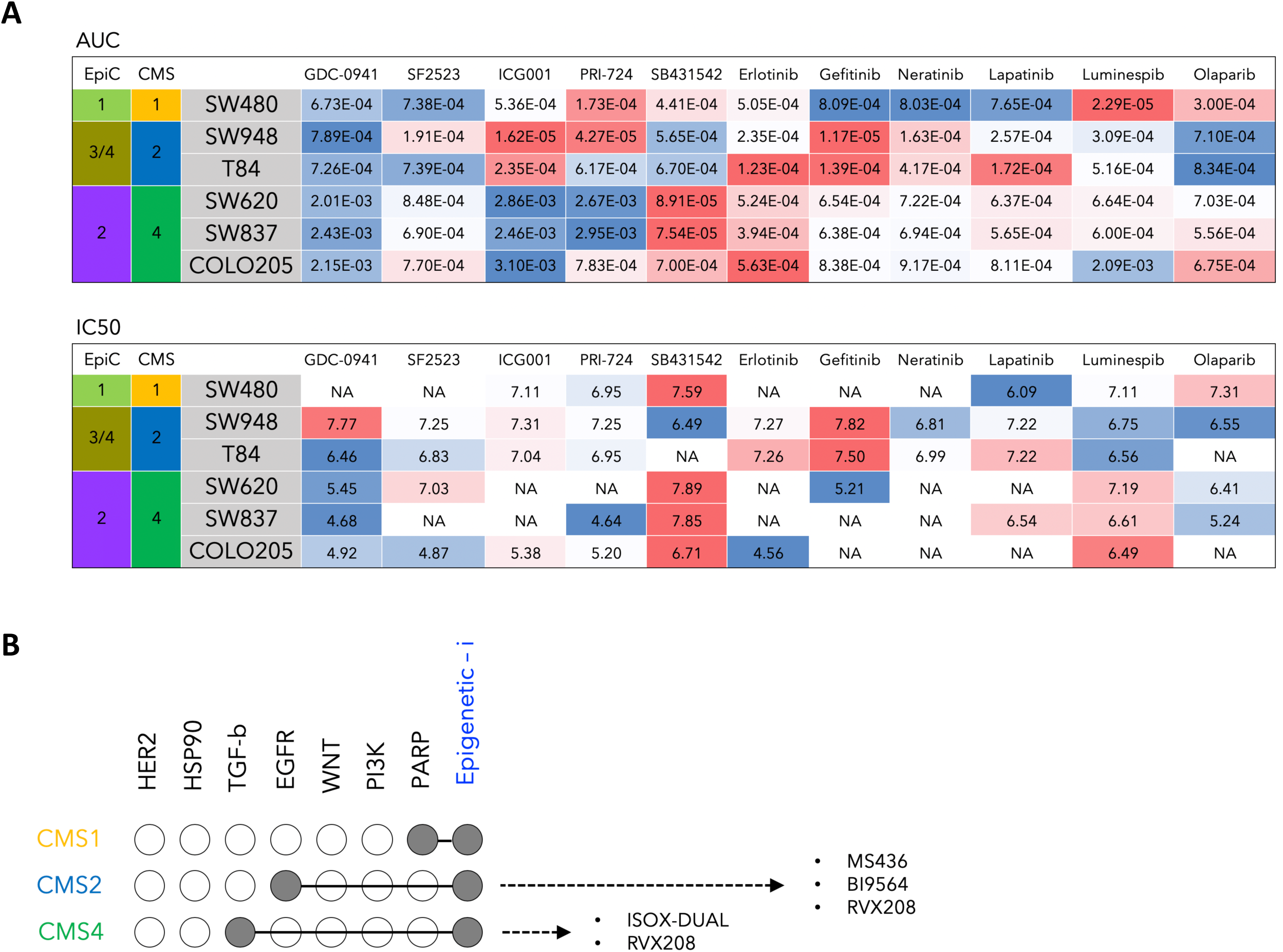

**Figure S14.**
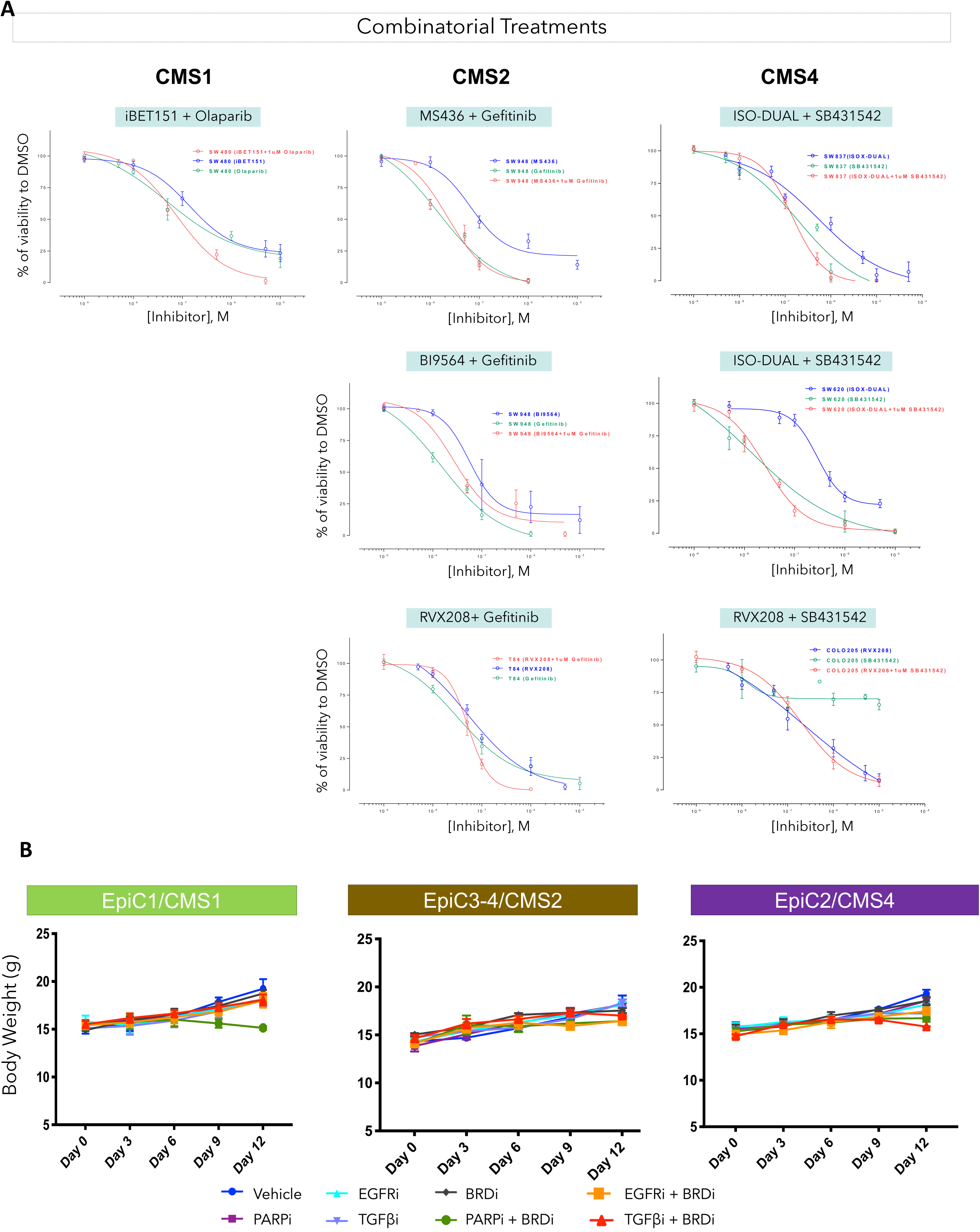

